# HILAMA: High-dimensional multi-omic mediation analysis with latent confounding

**DOI:** 10.1101/2023.09.15.557839

**Authors:** Xinbo Wang, Junyuan Liu, Sheng’en Shawn Hu, Zhonghua Liu, Hui Lu, Lin Liu, the Alzheimer’s Disease Neuroimaging Initiative

**Affiliations:** State Key Lab of Microbial Metabolism, Joint International Research Laboratory of Metabolic Developmental Sciences, Department of Bioinformatics and Biostatistics, School of Life Sciences and Biotechnology, Shanghai Jiao Tong University, Shanghai, China; SJTU-Yale Joint Center of Biostatistics and Data Science, National Center for Translational Medicine, Shanghai Jiao Tong University, Shanghai, China; School of Mathematical Sciences, Shanghai Jiao Tong University, Shanghai, China; Center for Public Health Genomics, University of Virginia, Charlottesville, VA, USA; Department of Biostatistics, Columbia University, New York, NY, USA; Shanghai Children’s Hospital, School of Medicine, Shanghai Jiao Tong University, Shanghai, China; Institute of Natural Sciences, MOE-LSC, CMA-Shanghai, Shanghai Jiao Tong University, Shanghai, China

## Abstract

**Motivation:** The increasingly available multi-omic datasets have posed both new opportunities and challenges to the development of quantitative methods for discovering novel mechanisms in biomedical research. One natural approach to analyzing such datasets is mediation analysis originated from the causal inference literature. Mediation analysis can help unravel the mechanisms through which exposure(s) exert the effect on outcome(s). However, existing methods fail to consider the case where (1) both exposures and mediators are potentially high-dimensional and (2) it is very likely that some important confounding variables are unmeasured or latent; both issues are quite common in practice. To the best of our knowledge, however, no methods have been developed to address these challenges with statistical guarantees.

**Results:** In this article, we propose a new method for HIgh-dimensional LAtent-confounding Mediation Analysis, abbreviated as “HILAMA”, that considers both high-dimensional exposures and mediators, and more importantly, the possible existence of latent confounding variables. HILAMA achieves false discovery rate (FDR) control under finite sample size for multiple mediation effect testing. The proposed method is evaluated through extensive simulation experiments, demonstrating its improved stability in FDR control and superior power in finite sample size compared to existing competitive methods. Furthermore, our method is applied to the proteomics-radiomics data from ADNI, identifying some key proteins and brain regions relating to Alzheimer’s disease. The results show that HILAMA can effectively control FDR and provide valid statistical inference for high dimensional mediation analysis with latent confounding variables.

**Availability:** The R package **HILAMA** is publicly available at https://github.com/Cinbo-Wang/HILAMA.

## 1 Introduction

The emergence of modern biotechnologies, such as high-throughput omics and multimodal neuroimaging, has led to the rapid accumulation of omic data at various levels, often including information on genomics, epigenomics, transcriptomics, proteomics, radiomics, and clinical records. It then becomes possible to thoroughly study complex diseases such as cancer and Alzheimer’s disease by integrating information from various scales [Subramanian et al., 2020, Kreitmaier et al., 2023]. For example, large collaborative consortia such as Alzheimer’s Disease Neuroimaging Initiative (ADNI) have collected information across all the levels mentioned above to help unravel the causal mechanisms of Alzheimer’s disease [Bao et al., 2023]. It is thus urgently needed to develop rigorous statistical methods for analyzing such datasets to reliably dissect the causal mechanisms [Tanay and Regev, 2017, Lv et al., 2021, Corander et al., 2022].

Such a problem falls into the category of causal mediation analysis [VanderWeele, 2015, Baron and Kenny, 1986], which can help disentangle the intermediate mechanisms between cause-effect pairs from observational datasets [Tobi et al., 2018, Liu et al., 2022, Clark-Boucher et al., 2023]. The classical mediation analysis can be traced back to Wright [1934] in 1930s, and then reverberated by Baron and Kenny [1986] in early 1980s based on regression techniques. Over the following decades, a vast literature has been engendered to put mediation analysis on a more rigorous ground both mathematically and conceptually [Robins and Greenland, 1992, Pearl, 2001, VanderWeele and Vansteelandt, 2014, Lindquist, 2012]. We refer interested readers to VanderWeele [2015] for a textbook-level introduction.

However, methods for mediation analyses with a single or a few exposures and/or mediators often cannot be directly scaled to address high-dimensional omic data. For instance, traditional hypothesis testing methods, such as Joint Significance Test (JST) [MacKinnon et al., 2002], Sobel’s method [Sobel, 2008], and bootstrap method [MacKinnon et al., 2007]], tend to be overly conservative particularly in genome-wide epigenetic studies [Barfield et al., 2017, Huang, 2019].

In response to the above challenge, recent years have seen a surge in the development of new methods for high-dimensional mediation analysis. These methods aim to explore the biological mechanisms derived from multi-omics data, as evidenced by studies such as Zeng et al. [2021] and Zhang et al. [2022a]. For instance, Zhang et al. [2016] and Gao et al. [2019] have focused on epigenetic studies with high-dimensional mediators and a continuous outcome. They have primarily employed (debiased) penalized linear regression and multiple testing procedures to formulate their methods. Derkach et al. [2019] have considered this similar problem by considering multiple latent variables as mediators that influence both the high dimensional biomarkers and the outcome. Moreover, Luo et al. [2020], Zhang et al. [2021] and Tian et al. [2022] have extended the analysis to include a survival outcome, in addition to high-dimensional mediators. These studies have contributed to the growing literature on exploring complex biological relationships. In a similar vein, Shao et al. [2021] has investigated high-dimensional exposures with a single mediator in an epigenetic study, utilizing a linear mixed-effect model.

However, there have been limited works studying both multivariate exposures and mediators. Zhang [2022] consider high-dimensional exposures and mediators through two different procedures; however, they require the mediators to be independent and mainly focus on mediator selection. Meanwhile, Zhao et al. [2022] develop a novel penalized principal component regression method that replaces the exposures with their principal components in a lower dimension. This approach, however, lacks causal interpretation. More importantly, most high-dimensional mediation analyses approaches make the untestable assumption of no latent confounding, which is highly problematic in multi-omic biological studies due to the prevalence of non-randomized study designs. If hidden confounding cannot be eliminated, these methods may be influenced by spurious correlations and result in an inflated False Discover Rate (FDR). Recently, several works have addressed this issue by considering latent confounding in high-dimensional linear models, which involve estimating overall causal effects under the Latent Structural Equation Modeling (LSEM) framework [Chernozhukov et al., 2017, Ćevid et al., 2020, Guo et al., 2022, Bing et al., 2022a,b]. Specifically, Sun et al. [2022] are the first to address the large-scale hypothesis testing problem in the high-dimensional confounded linear model. They achieve FDR control under finite sample size by introducing a decorrelating transformation before the debiasing step.

Inspired by the aforementioned works on high-dimensional linear regression under latent confounding [Chernozhukov et al., 2017, Ćevid et al., 2020, Guo et al., 2022, Bing et al., 2022a,b, Sun et al., 2022], we propose a novel method called HILAMA, which stands for HIgh-dimensional LAtent-confounding Mediation Analysis. HILAMA addresses two critical challenges in applying mediation analysis (or any causal inference method) to multi-omics studies: (1) accommodating both high-dimensional exposures and mediators, and (2) handling latent confounding. In contrast to competing methods [Baron and Kenny, 1986, Zhang et al., 2016, Gao et al., 2019, Schaid et al., 2022, Zhao et al., 2022], our method maintains control over FDR at the nominal level for multiple mediation effect testing, even in the presence of latent confounders. Now, we briefly sketch the essential components of our method. First, we employ the Decorrelate & Debias method in Sun et al. [2022] to obtain p-values for each individual exposure and mediator’s effect on the outcome. Second, to estimate the effect matrix of exposures on mediators, we employ a column-wise regression strategy, again incorporating the Decorrelate & Debias method [Sun et al., 2022]. To handle large and high-dimensional datasets, we utilize parallel computing in this step. Third, we apply the Min-Screen procedure in Djordjilovi et al. [2019] to eliminate non-rejected hypotheses, retaining only the *K* most significant pairs for the final stage of multiple testing. Lastly, we compute p-values for all *K* pairs using the JST method [MacKinnon et al., 2002], employing a data-dependent threshold determined by the Benjamini-Hochberg (BH) procedure [Benjamini and Hochberg, 1995] to maintain FDR at the nominal level *α*. We conduct extensive simulations to evaluate our method’s performance, which demonstrates effective FDR control across various finite sample sizes, surpassing the capabilities of most other methods. Furthermore, we apply HILAMA to a proteomics-radiomics dataset from the ADNI database (adni.loni.usc.edu) and identify key proteins and brain regions associated with learning, memory, and recognition impairments in Alzheimer’s disease and cognitive impairment.

The rest of this article is organized as follows. In Section 2, we first describe our model and then introduce the HILAMA procedure under the Linear Structural Equation Models with highdimensional exposures, high-dimensional mediators, continuous outcome, and latent confounders setting. In Section 3, we evaluate the FDR and Power performance of HILAMA across a wide range of simulations. In Section 4, we apply the HILAMA to a proteomics-radiomics data of Alzheimer’s disease from ADNI. The paper is concluded with a discussion in Section 5. Technical details, some tables and figures are collected in the online Supplementary Materials.

## 2 Methods

To describe the HILAMA methodology, we first need to briefly review mediation analyses. Mediation analyses are frequently utilized to disentangle the underlying causal mechanism between two sets of variables, the exposures and outcomes, exerted by a third set of variables, the mediators. The overall causal effects can be decomposed into direct effects from exposures to outcomes, bypassing mediators, and indirect effects via mediators. To be more precise, we ground our discussion under the Linear Structural Equation (LSE) framework, that models the causal mechanisms among *p*-dimensional exposures ***X***_*i*_ = (*X*_*i*1_, ⋯, *X*_*ip*_)^⊤^ ∈ ℝ^*p*^, *q*-dimensional mediators ***M***_*i*_ = (*M*_*i*1_, ⋯, *M*_*iq*_)^⊤^ ∈ ℝ^*q*^, a scalar outcome *Y*_*i*_ ∈ ℝ, and latent confounders ***H***_*i*_ = (*H*_*i*1_, ⋯, *H*_*is*_)^⊤^ ∈ ℝ^*s*^ (e.g., batch effects, disease subtypes, and lifestyle factors) as follows:

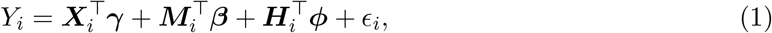

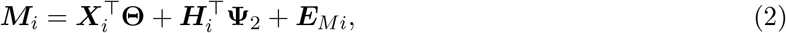

where *ϵ*_*i*_ and ***E***_*Mi*_ are the noise terms that are independent of ***X***_*i*_, ***M***_*i*_ and ***H***_*i*_. In the outcome model (1), ***γ*** = (*γ*_1_, ⋯, *γ*_*p*_)^⊤^ ∈ ℝ^*p*^ is the direct effect vector of the exposures ***X***_*i*_ on the outcome *Y*_*i*_, and ***β*** = (*β*_1_, ⋯, *β*_*q*_)^⊤^ ∈ ℝ^*q*^ represents the effect vector of the mediators ***M***_*i*_ to the outcome *Y*_*i*_ after adjusting for the latent confounders ***H***_*i*_. ***ϕ*** ∈ ℝ^*s*^ is the parameter vector that relates latent confounders ***H***_*i*_ to the outcome *Y*_*i*_. Here, we allow *p* and *q* to be larger than the sample size *n*, while *s* ≤ *p* + *q*. The primary objective of our study is to identify the active direct/indirect effects (*γ*_*k*_, *θ*_*kl*_*β*_*l*_, *k* ∈ [*p*], *l* ∈ [*q*]) from *p/pq* possible paths, as shown in Figure 1.

**Fig. 1.**
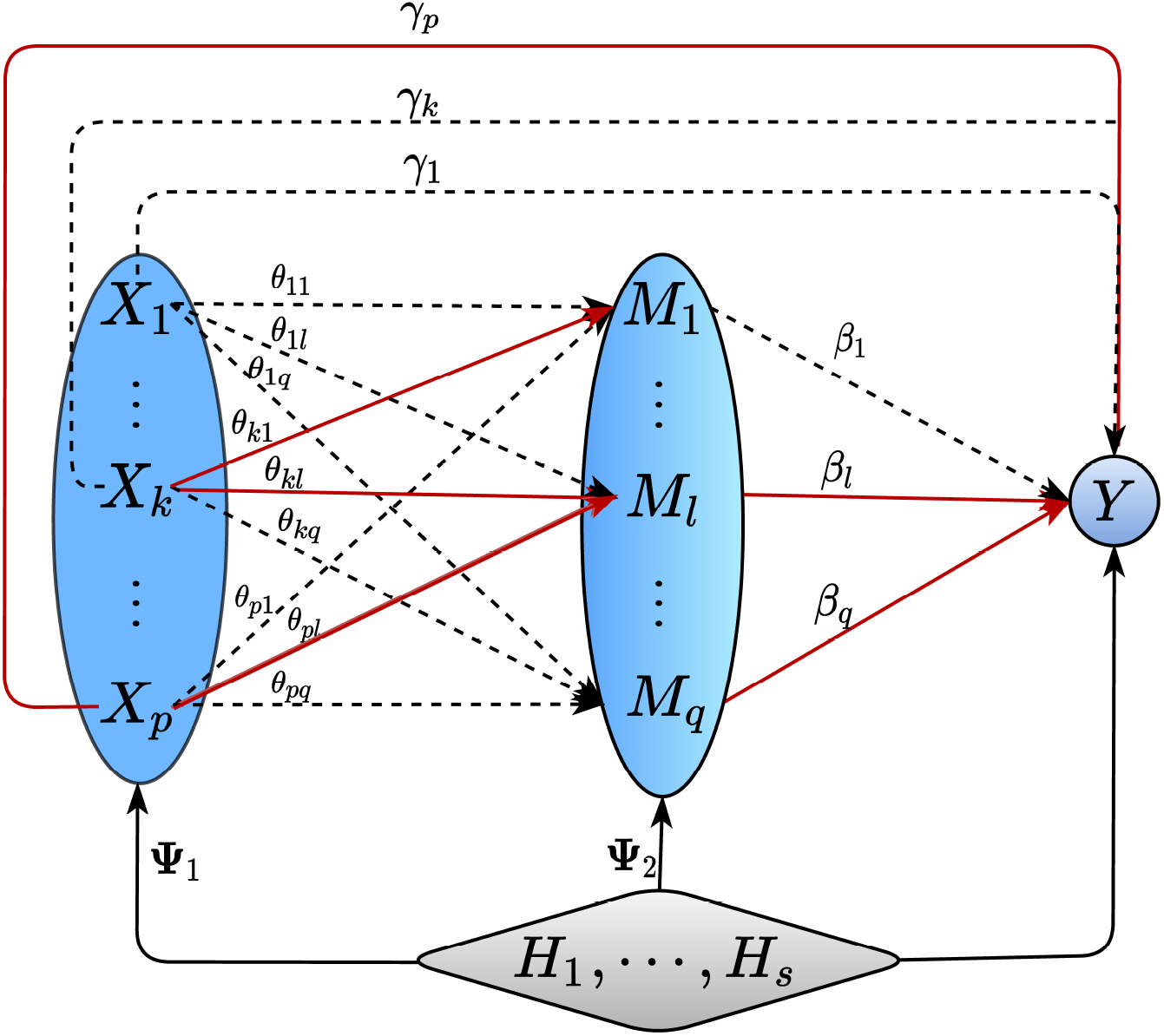
Causal Diagram for the setup considered in this paper. ***X*** = (*X*_1_, ⋯, *X*_*p*_)^⊤^: exposures of dimension *p* (protein expression data in our real-data applications); ***M*** = (*M*_1_, ⋯, *M*_*q*_)^⊤^: mediators of dimension *q* (imaging data in our real-data applications); *Y* : a real-valued outcome (clinical outcome or phenome data in our real-data applications); ***H*** = (*H*_1_, ⋯, *H*_*s*_)^⊤^: latent/unmeasured confounders of dimension *s* (e.g. mis-measured clinical data, epigenetic information, etc.). Solid (red) lines indicate non-null effects, while dotted lines indicate null effects.

In the mediator model (2), the matrix **Θ** = (***θ***_1_, ⋯, ***θ***_*q*_) = (*θ*_*kl*_) ∈ ℝ^*p*×*q*^ represents the regression coefficients of exposures on mediators and *θ*_*kl*_ represents the effect of exposure *X*_*ik*_ on mediator *M*_*il*_ after adjusting for the effect of latent confounders ***H***_*i*_. **Ψ**_2_ ∈ ℝ^*s*×*q*^ can be interpreted as the confounding effect of the latent confounders ***H***_*i*_ on mediators ***M***_*i*_. The mediation model (1) – (2) adopted here is similar to those proposed by Schaid et al. [2022] and Zhao et al. [2022], which were among the first works to consider both multivariate exposures and mediators. However, we incorporate the latent confounders into our high-dimensional mediation analysis, which is a novel approach in the field. Furthermore, unlike the approach used in Zhao et al. [2022], our mediation analysis is directly based on ***X***_*i*_, instead of a transformation 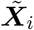 of the original vector ***X***_*i*_.

As mentioned, the causal parameters of interest in mediation analyses are mainly the (average) natural direct and indirect effects. When the ignorability assumption (explained in Supplementary Materials) holds, the natural direct effect of exposure *k* on outcome, denoted by NDE_*k*_, and the nature indirect effect of exposure *k* on outcome, denoted by NIE_*k*_, can be expressed as [Robins and Greenland, 1992, Pearl, 2001]

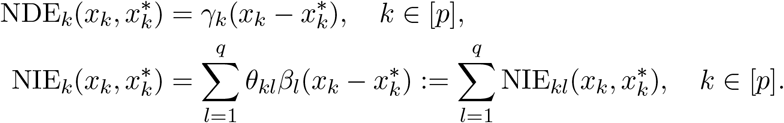

When *x*_*k*_ and 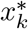 defer by one unit, NDE_*k*_(1) = *γ*_*k*_, the regression coefficient between ***X*** and *Y* in model (1), and NIE_*kl*_(1) = *θ*_*kl*_*β*_*l*_, the product of the regression coefficient between ***X***_*k*_ and ***M***_*l*_ in model (2) and the regression coefficient between ***M***_*l*_ and *Y* in model (1). For their derivation, see Supplementary Materials.

However, when latent confounders exist (i.e. dim(***H***) *>* 0), neither NDEs nor NIEs are identifiable without making additional assumptions, which are generally based by domain-specific knowledge. To identify the true parameter ***γ, β*** and ***θ***_*l*_ in the confounded linear model (1) and (2), it is necessary to make additional assumptions among the observed variables (***X***_*i*_, ***M***_*i*_) and the latent confounders ***H***_*i*_. Based on the works of Wang et al. [2017], Ćevid et al. [2020], Guo et al. [2022], Sun et al. [2022], a factor model is specified to characterize the relation between ***X***_*i*_ and ***H***_*i*_:

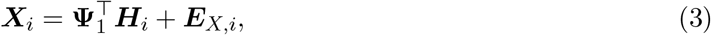

where Cov(***H***_*i*_, ***E***_*X,i*_) = 0 and the random variable ***E***_*X,i*_ ∈ ℝ^*p*^ represents the unconfounded components of ***X***_*i*_. Moreover, to accurately identify the true signals and effectively remove the confounding effects, we impose a spiked singular value condition on the covariance between the exposures and mediators, as shown in Figure 6. Specifically, we require 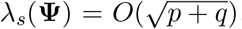, where **Ψ** = (**Ψ**_1_, **Ψ**_2_) ∈ ℝ^*s*×(*p*+*q*)^. Our approach is particularly effective in scenarios where the confounding effect is dense, i.e., many observed variables in ***X*** ∈ ℝ^*p*^ and ***M*** ∈ ℝ^*q*^ are simultaneously influenced by the latent confounders ***H*** ∈ ℝ^*s*^.

Our goal is to identify the path-specific indirect effect *θ*_*kl*_*β*_*l*_ (corresponding to the path *X*_*k*_ → *M*_*l*_ → *Y*) from the total *p* · *q* possible paths which corresponds to the following multiple hypothesis testing problem:

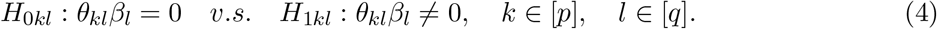

Here, we propose a novel framework called HILAMA to address the problem (4). The framework identifies the true paths with nonzero indirect effects and controls the finite-sample FDR. It involves four major steps explained in detail below (as shown in Figure 2).

**Fig. 2.**
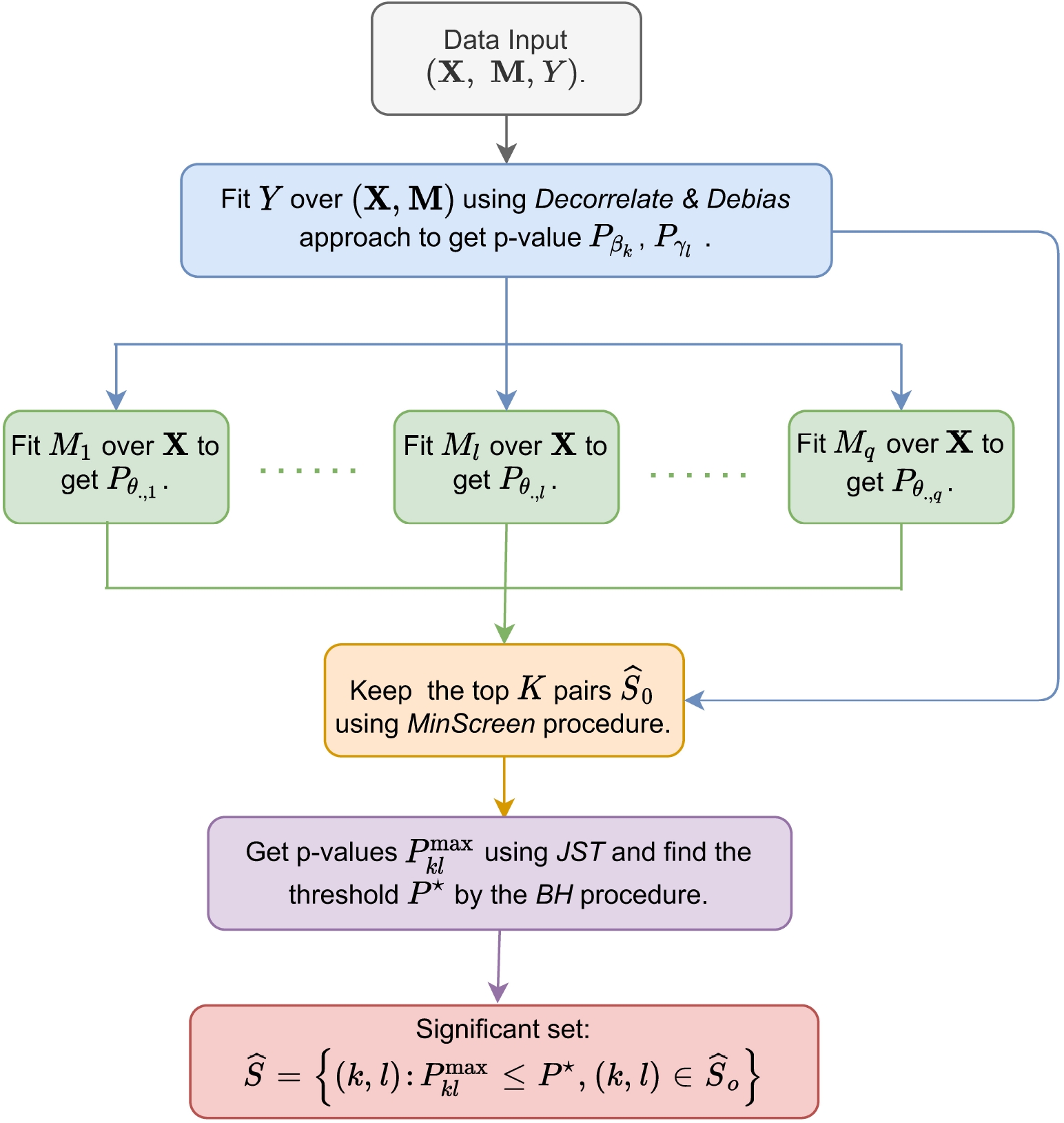
Flowchart of HILAMA. First, we regress outcome *Y* over mediators ***M*** and exposures ***X*** using the *Decorrelate* & *Debias* approach to obtain the debiased p-values for the parameter ***β***. Second, we similarly regress each mediator *M*_*l*_ over the expsoures ***X*** separately to obtain debiased p-values for the parameter **Θ**. Third, we employ the *MinScreen* procedure to select a subset of *K* pairs for subsequent multiple testing. Finally, we calculate p-values for the mediation effect using the *JST* method and choose a p-value threshold using the *BH* procedure based on a pre-specified FDR level, where a set of pairs is considered significant if the p-value falls below this threshold.

First, for the outcome model in equation (2), we utilize the *Decorrelate* & *Debias* approach presented in Sun et al. [2022] to carry out inference on the regression parameters ***γ*** and ***β***. By applying this method, we obtain the double debiased estimator 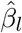 and 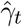 These estimators, after appropriate rescaling, asymptotically converge to centered Gaussian distributions: 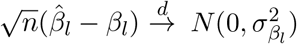 and 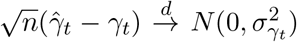 individually under some mild conditions. We denote the corresponding variance estimators as 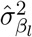 and 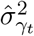 Then p-values can be computed as follows:

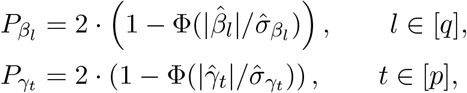

where Φ(·) denotes the cumulative distribution function of the standard normal distribution *N* (0, 1).

Second, to estimate each column of the parameter matrix **Θ** in the multi-response mediation model defined in equation (2), we employ a column-wise regression strategy. For each sub-regression problem, we utilize the *Decorrelate* & *Debias* approach [Sun et al., 2022]. To handle the computational challenges posed by large-scale datasets in multi-omics studies, we leverage parallel computing techniques to accelerate the computation process. This allows us to efficiently calculate the point and variance estimators for the coefficients *θ*_*kl*_(*k* ∈ [*p*] and *l* ∈ [*q*]), which are denoted as 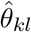 and 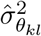, respectively. To assess the statistical significance of the coefficient estimates, we calculate the p-values using the formula:

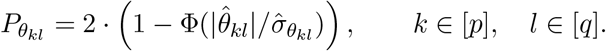

where Φ denotes the cumulative distribution function of the standard normal distribution.

Third, we employ the MinScreen procedure [Djordjilovi et al., 2019] to screen the total *p* · *q* possible causal paths. The screened causal paths by MinScreen are defined as the top *K* significant paths: 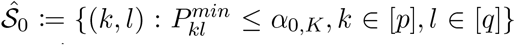 where 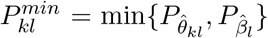 and *α*_0_ is chosen such that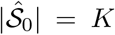 This preliminary step eliminates the least promising causal paths before calculating the final p-value for *H*_0*kl*_. By doing so, it effectively reduces the computational burden in the subsequent multiple testing phase.

Lastly, we apply the joint significance test (JST), also known as the MaxP test [MacKinnon et al., 2002], to obtain the p-value for the null hypothesis *H*_0*kl*_ : *θ*_*kl*_*β*_*l*_ = 0 which tests for no indirect effect, for 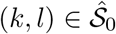 The p-values for JST are defined as

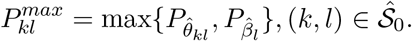

We then sort the JST p-values and denote them as *p*_(*i*)_, *i* = 1, ⋯, *K*, the notation for order statistics by convention. To protect the FDR at the nominal level *α*, we follow the BH procedure [Benjamini and Hochberg, 1995] to find the data-driven p-value rejection threshold *P*\* as

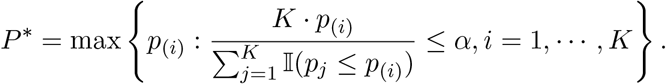

Finally, we define the set containing statistically significant non-zero path-specific effects as 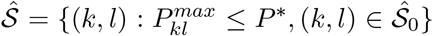 In this article, we evaluate HILAMA and other competitors by FDR and Power, which are defined as follows:

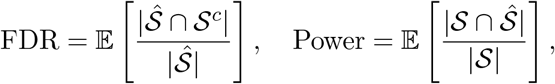

where 𝒮 = {(*k, l*) : *θ*_*kl*_*β*_*l*_ ≠ 0, *k* ∈ [*p*], *l* ∈ [*q*]} represents the true non-zero effect path-specific set and 𝒮^*c*^ = {(*k, l*) : *θ*_*kl*_*β*_*l*_ = 0, *k* ∈ [*p*], *l* ∈ [*q*]} represents the zero effect path set.

## 3 Simulation Studies

In this section, we assess if HILAMA is capable of controlling the FDR with sufficient power across a wide range of simulation settings. The performance is compared against various other approaches.

As a baseline benchmark, we employ the univariate Baron & Kenny method [Baron and Kenny, 1986] (abbreviated as BK) for every possible individual exposure-mediator pair, using the R package **mediation**. We also consider methods that only allow a *univariate* exposure and high-dimensional mediators, including HIMA [Zhang et al., 2016] and HDMA [Gao et al., 2019]. For these two methods, we directly use their corresponding R packages **HIMA** and **HDMA**, analyze every individual exposure, and then aggregate the results. Finally, we compare two penalized methods developed for multiple exposures and mediators. Specifically, for the method “mvregmed” [Schaid et al., 2022], we apply the R package **regmed**. While for the method developed by Zhao et al. [2022] (abbreviated as ZY), we implement their penalized regression algorithm and omit the dimension reduction step for comparison. Here, we only compare the two penalized methods in the low dimensional setup in **simulation 2** introduced below due to their slow running time (See Figure 4d).

We first generate the exposure data ***X***_*i*_(*i* = 1, ⋯, *n*) according to model (3). The latent confounders ***H***_*i*_ ∈ ℝ^*s*^ and the elements of confounding matrix **Ψ** ∈ ℝ^*s*×*p*^ are independently drawn from the standard normal distribution. The unconfounded components ***E***_*X,i*_ are drawn from *N*_*p*_(0, **Σ**_*E*_), where **Σ**_*E,kl*_ = *κ*^|*k*−*l*|^(*k, l* ∈ [*p*]). The parameter *κ* controls the strength of correlation among exposures, and it takes values in the range [0, 1).

Similarly, we generate the the mediator data ***M***_*i*_(*i* = 1, ⋯, *n*) according to model (2). The noise term ***E***_*M,i*_ are drawn from *N*_*q*_(0, ***I***) and the confounding effect matrix **Ψ**_2,*kl*_(*k* ∈ [*s*], *l*∈ [*q*]) are drawn from *ξ* · *N* (*η*, 1), where *ξ* is a Rademacher random variable, i.e. 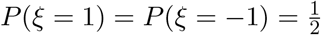 Then, for the signal coefficient matrix **Θ** ∈ ℝ^*p*×*q*^, we randomly choose *p* · *r*_*p*_ rows having non-zero elements, and choose *δ* non-zero elements seperately in each of these rows, where *δ* follows uniform distribution on {5, 6, ⋯, 20}. The non-zero elements in **Θ** follow uniform distribution on [−1.5, −0.5] ∪ [0.5, 1.5].

Finally, we generate the outcome data *Y*_*i*_(*i* = 1, ⋯, *n*) according to model (1). The coefficients ***γ*** are randomly selected with *p*·*r*_*p*_ non-zero elements following a uniform distribution on [−1.5, −0.5]∪ [0.5, 1.5]. As for the coefficients ***β***, we choose *q*·*r*_*q*_ non-zero elements following a uniform distribution on [−1.5, −0.5] ∪ [0.5, 1.5]. To determine the active location in ***β***, we define 𝒜^*c*^ as the set of columns in **Θ** with zero elements (∥Θ_.*l*_∥_1_ = 0), and 𝒜 as the set of columns in **Θ** with non-zero elements (∥Θ_.*l*_∥_1_ ≠ 0). From 𝒜^*c*^, we randomly choose *s*_01_ elements with equal probability, where *s*_01_ = min{0.2 · *q* · *r*_*q*_, |𝒜^*c*^|}. While from 𝒜, we randomly choose *s*_11_ elements with unequal probability, where *s*_11_ = *q* · *r*_*q*_ − *s*_01_. The selection probability of *l*∈𝒜 is determined by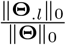which represents the proportion of non-zero elements in column *l* relative to all non-zero elements in **Θ**. The confounding effects ***ϕ*** are drawn from *N*_*s*_(*η, I*) · *ξ*, and the noise terms *ϵ*_*Y,i*_ are drawn from *N* (0, 1).

For all the simulations below, we fix the sparsity proportions as *r*_*p*_ = *r*_*q*_ = 0.1 and the dimension of latent confounders *s* = 3. Additionally, we set the nominal FDR level at the *α* = 0.1 and all the simulation results are averaged over 50 Monte Carlo replications.

### Simulation 1

In the first simulation, we test the stability of our model under various scenarios. We evaluate the impact of changes in sample size (*n* ∈ {200, 400}), exposure dimension (*p* ∈ {100, 200, 400}), mediator dimension (*q* ∈ {50, 100, 200}), correlation size among exposures (*κ* ∈ {0.4, 0.8}), and magnitude of latent effects (*η* ∈ {0.5, 1.5}). For the total 72 different settings, we present the average value of empirical FDR and Power in Figure S1a and Figure S1b.

For simplicity, we only present scenarios for *p* = 400 in Figure 3. From Figure 3a, only HILAMA controls the FDR at the nominal level *α* = 0.1 in all scenarios, whereas the other three methods all fail to do so. The reasons for their lack of control are due to their failure to correct for the effect of latent confounding, and their inability to accommodate high-dimensional exposure and mediator settings. Turning to the power, by reading Figure 3b, we can easily see that HILAMA achieves the highest and most stable power close to 1 in large sample sizes (*n* = 400). However, in smaller sample sizes (*n* = 200), the statistical power decreases as correlation coefficient *κ* or confounding effect *η* increases. The power of HILAMA is generally unaffected by the above parameters in larger samples. The powers of the other three methods, on the other hand, are essentially meaningless since their FDRs are all close to 1. Moreover, the point estimates of mediation effects output by HILAMA has much smaller bias than the other competing methods, represented as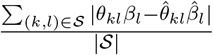 (see Figure S1c).

**Fig. 3.**
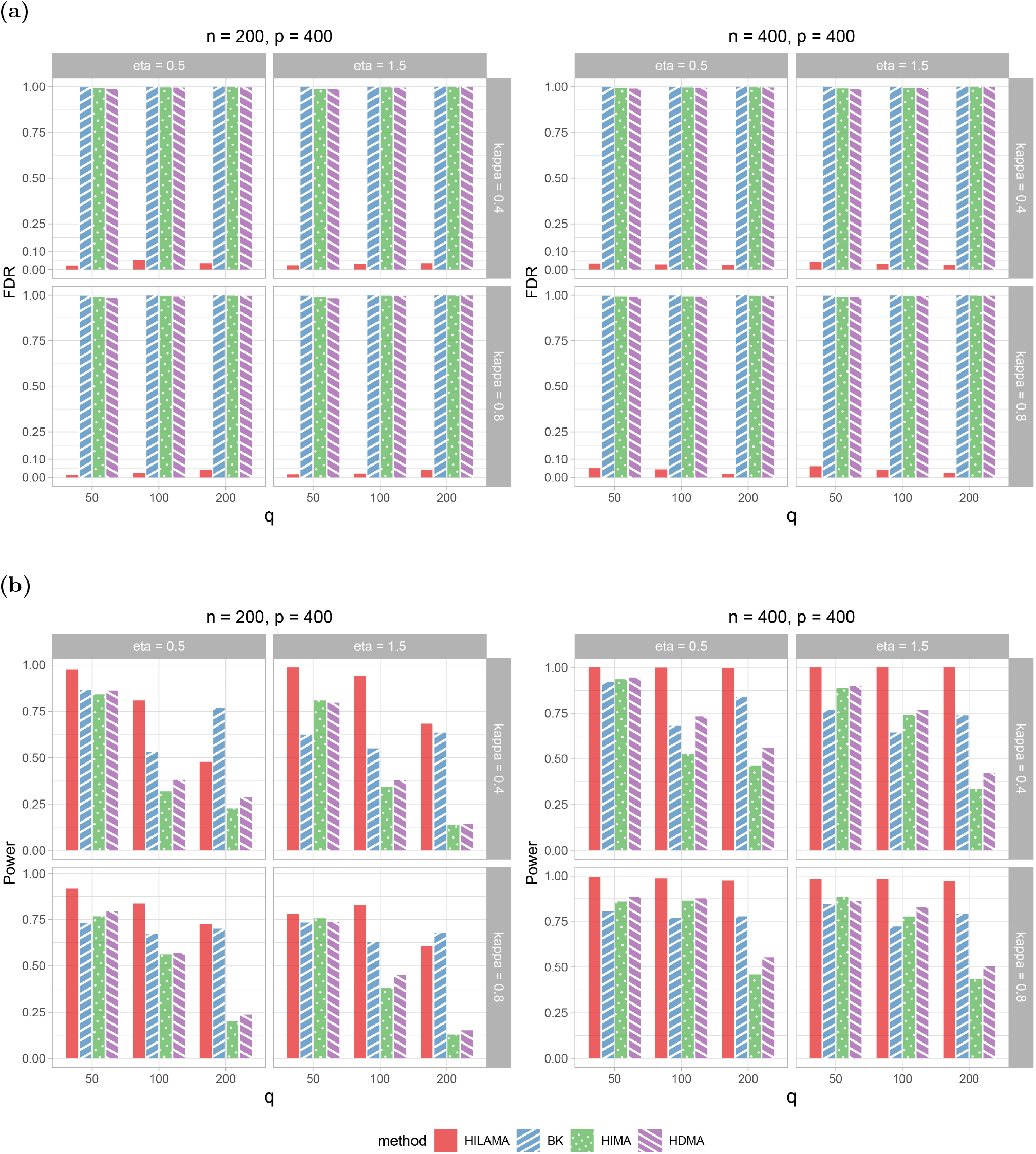
Comparison results of (a) Empirical False Discovery Rate (FDR) and (b) Empirical Power for different methods in **simulation 1** across 72 scenarios. *eta* represents latent effect and *kappa* represents the correlation size among exposure. All results are averaged over 50 replications.

### Simulation 2

In the second simulation, we assess how the denseness of latent confounding impacts the performance of HILAMA. We measure the denseness of latent confounding as the proportion (1− *r*_*h*_) of zero entries in each row of **Ψ**_1_ and **Ψ**_2_. If *r*_*h*_ = 0, then **Ψ**_1_ = **Ψ**_2_ = 0, amounting to no latent confounding; whereas if *r*_*h*_ = 1, all exposures and mediators are confounded by latent confounders, as depicted in simulation 1. Here, we vary only *r*_*h*_ ∈ {0, 0.1, 0.2, ⋯, ⋯ 0.9, 1} while holding *n* = 400, *p* = *q* = 100, *κ* = 0.6, *η* = 1 and compare HILAMA with the two aforementioned penalized methods. To compare our p-value based method with the penalized methods mvregmed and ZY, we assume that the actual number of active pairs is known. We select the top *K* pairs that control the FDR at the level 0.1 and compare their power. If the FDR cannot be controlled at the 0.1 level, we choose the cut-off point associated with the lowest FDR and calculate the corresponding power.

Figure 4a indicates that HILAMA does not manage the FDR at the 0.1 level under the low-dimensional setting when only a few observed variables are confounded by latent confounders. Additionally, the power of HILAMA is highly sensitive when a few observed variables are confounded by latent confounders from Figure 4b, although it tends to stabilize as the confounding density increases. However, the mvregmed method does not control the FDR across all situations, even in the absence of latent confounding. Figure 4c demonstrates that HILAMA again has the minimum mean bias compared to the other two competing methods, mvregmed and ZY. Specifically, Figure 4d shows that although the ZY method boasts the best FDR control and power performance, it takes hundreds of times longer to compute than HILAMA, even in this low-dimensional setting.

**Fig. 4.**
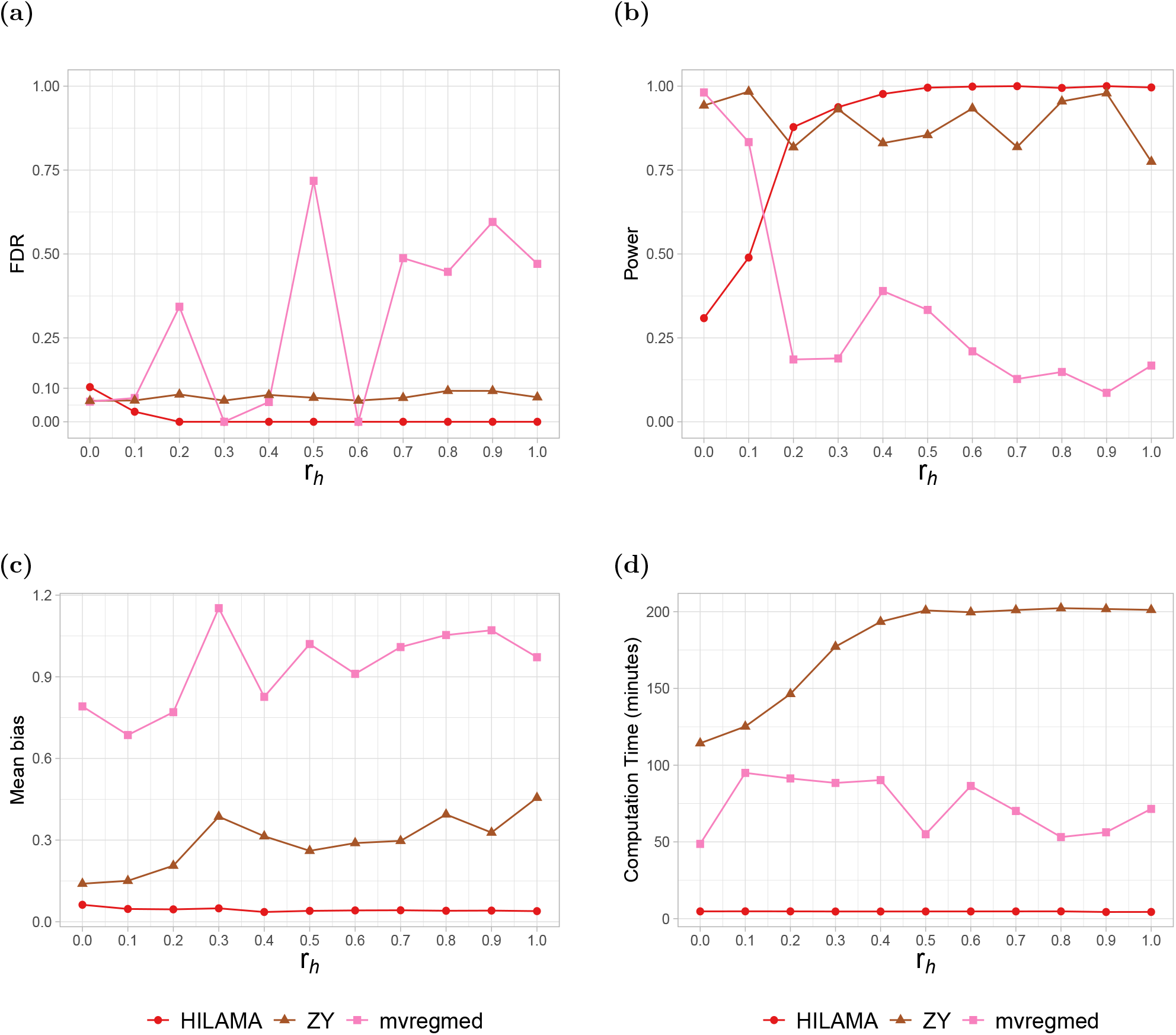
Comparison results of (a) Empirical FDR, (b) Empirical Power, (c) Mean bias and (d) Computation time (minutes) for different methods in **simulation 2 with low dimensional setting** across varied confounding density *r*_*h*_ while fixing *n* = 400, *p* = *q* = 100, *κ* = 0.6, *η* = 1, *r*_*p*_ = *r*_*pq*_ = 0.1, *s* = 3. All results are averaged over 50 replications.

Additionally, we consider *n* = 300, *p* = 200, *q* = 100, while all other components of the data generating distribution are held constant and compare HILAMA with the above three p-value based methods. Figure 5a and Figure 5b display the FDR and power of the four p-value-based methods at the nominal FDR level of 0.1. HILAMA demonstrates effective FDR control in all cases, including under no latent confounding, whereas the other three methods are unable to control the FDR in any setting. HILAMA sustains a stable power near 1 in most scenarios, except when *r*_*h*_ = 0. Meanwhile, the powers of HIMA and HDMA have a downward trend as *r*_*h*_ increases. Figure 5c additionally explores the average bias of true mediation effect and it is obvious that HILAMA attains the minimum mean bias across all the scenarios, while all the other other methods show an increasing trend in mean bias as *r*_*h*_ increases.

**Fig. 5.**
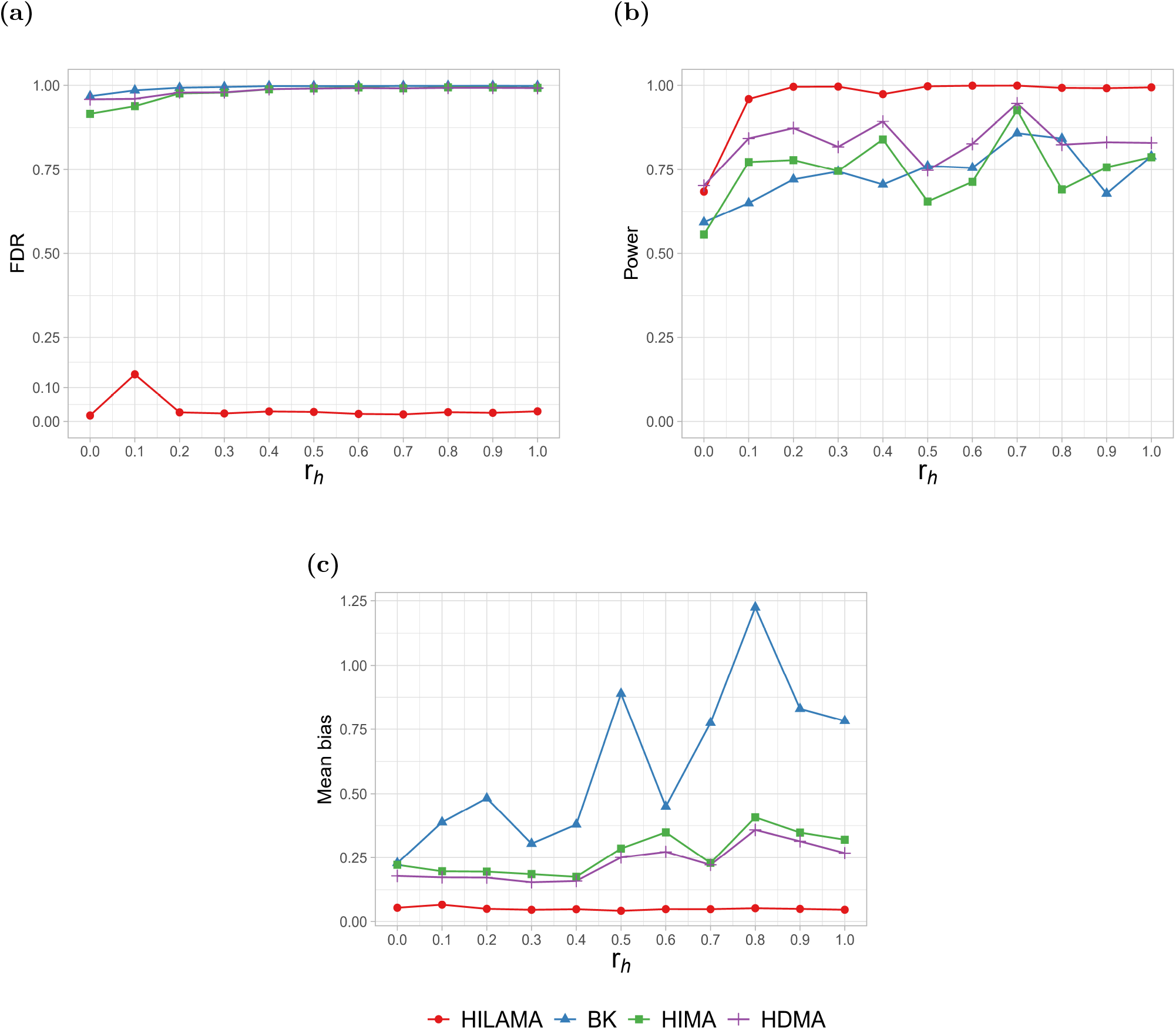
Comparison results of (a) Empirical FDR, (b) Empirical Power and (c) Mean bias for different methods in **simulation 2 with high dimensional setting** across varied confounding density *r*_*h*_ while fixing *n* = 300, *p* = 200, *q* = 100, *κ* = 0.6, *η* = 1, *r*_*p*_ = *r*_*pq*_ = 0.1, *s* = 3. All results are averaged over 50 replications under the nominal FDR level of 0.1.

### Simulation 3

In the third simulation, our aim is to investigate the impact of signal strength on HILAMA. Specifically, the data generation process repeats that of the previous simulations, with the exception that the non-zero components of **Θ** follow the distribution of *ξ* · *N*(*ρ*, 1) and the non-zero elements of ***β*** follow *N*(*ρ*, 0.1). We vary the signal strength *ρ* ∈ {0.1, 0.2, ⋯, 1.4, 1.5}while maintaining *n* = 400, *p* = 300, *q* = 100, *κ* = 0.6, and *η* = 1.

According to Figure S2a, HILAMA can control the FDR to the nominal level of 0.1, whereas the other three methods fail to do so. Notably, from Figure S2b we can see that the power of HILAMA is restricted when *ρ* - the signal strength is weak, and the power increases as *ρ* gets stronger, before stabilizing around 1 when *ρ* ≥1. Similar to these above findings, the mean bias of HILAMA is the smallest among all the methods being compared, as shown in Figure S2c.

## 4 An application to the Proteomics-Radiomics Study of AD

In this final section, we apply HILAMA to a real multi-omic dataset collected by the ADNI. Before delving into the details, we emphasize that this data analysis should be viewed as at most exploratory rather than confirmatory nature. It is highly likely that the linearity assumption imposed in the Structural Equation Model may not be a good approximation of the reality.

Alzheimer’s disease (AD) is an irreversible and complex neurological disease that affects millions of individuals worldwide. Currently, approximately 6.7 million Americans aged 65 years and older live with AD, and this number is projected to dramatically increase to 13.8 million by the year 2060 [AD2, 2023]. AD is characterized by progressive memory loss and other cognitive impairments resulting from the accumulation of amyloid-*β*(A*β*) and tau proteins in the brain, leading to neurodegenerative symptoms [Chen and Xia, 2020].

Unfortunately, there is currently no effective treatment for AD, underscoring the significance of early diagnosis and comprehending the disease’s pathogenesis. Therefore, it is crucial to develop effective interventions to prevent, slow down, or even cure this disease through biomedical research. With this in mind, the Alzheimer’s Disease Neuroimaging Initiative (ADNI, adni.loni.usc.edu) was established in 2003. Its primary goals are to develop biomarkers for AD, enhance the understanding of its pathophysiology, and improve early detection using various modalities such as magnetic resonance imaging (MRI), positron emission tomography (PET), functional magnetic resonance imaging (fMRI), as well as clinical and neuropsychological assessments.

In this section, we utilize the HILAMA approach to examine the connection between proteins in the cerebrospinal fluid (CSF), whole-brain atrophy, and cognitive behavior. Our aim is to identify critical biological pathways associated with AD by utilizing data from the ADNI database. The CSF proteomics data is acquired using a highly specific and sensitive technique called targeted liquid chromatography multiple reaction monitoring mass spectrometry (LC/MS-MRM), resulting a list of 142 annotated proteins derived from 320 peptides. Additionally, the brain imaging data is obtained through anatomical magnetic resonance imaging (MRI), and volumetric measurements are extracted from 145 brain regions-of-interest (ROI) [Doshi et al., 2016]. To assess the relationship between the aforementioned variables and cognitive function, we consider the composite memory score as the response. This score is measured using the ADNI neuropsychological battery, with higher scores indicating better cognitive function. In our model, we treat the 142 proteins as exposures (***X***), the 145 brain regions as mediators (***M***), and the memory score as the outcome (***Y***). For this study, we focus on a total of 287 subjects who have both proteomics and imaging data available. These subjects consist of 86 cognitively normal individuals (CN), 135 patients with mild cognitive impairment (MCI), and 66 AD patients. To account for potential confounding effects, we include covariates such as age, gender (Male = 1, Female = 2), years of education, and disease type (CN = 1, MCI = 2, AD = 3). For more detailed information on these baseline covariates, please refer to Table 1.

**Table 1.**
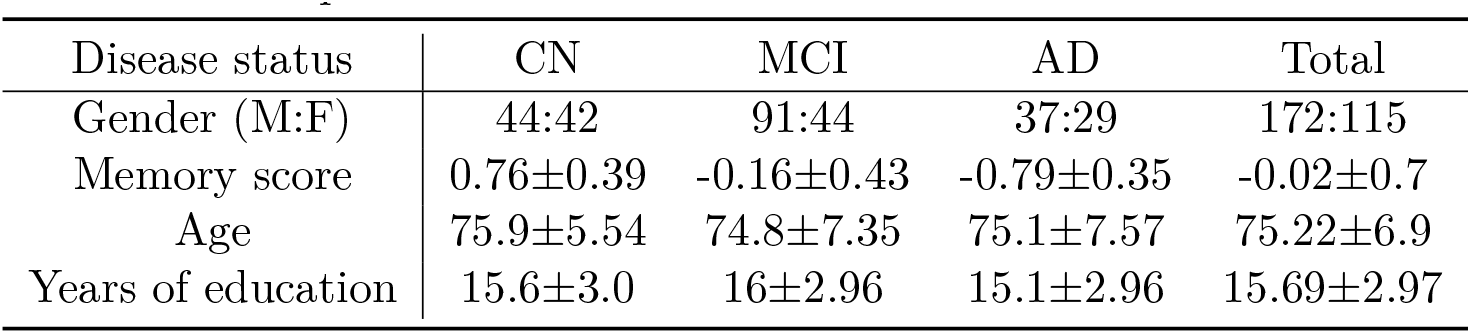
Frequencies and descriptive statistics (mean ± standard deviation) for demographic and clinical variables in the sample.

| Disease status | CN | MCI | AD | Total |
| --- | --- | --- | --- | --- |
| Gender (M:F) | 44:42 | 91:44 | 37:29 | 172:115 |
| Memory score | 0.76±0.39 | -0.16±0.43 | -0.79±0.35 | -0.02±0.7 |
| Age | 75.9±5.54 | 74.8±7.35 | 75.1±7.57 | 75.22±6.9 |
| Years of education | 15.6±3.0 | 16±2.96 | 15.1±2.96 | 15.69±2.97 |

Prior to conducting the mediation analysis, we impute some volumetric measures recorded as zero with the corresponding median value, and then apply a log-transformation to achieve a more normal distribution. Subsequently, we standardize both the protein data and MRI data to have a mean of zero and a standard deviation of one, while only centering the outcome cognitive score to have a mean of zero. In Figure 6, we visualize the singular values of the protein and MRI data, allowing us to assess the potential presence of latent confounders. By examining Figure 6a and Figure 6b, we observe the presence of two significantly larger singular values in the protein data and one in the MRI data. This finding suggests a distinct spiked structure, indicating the possible presence of latent confounders as depicted in the models (2) and (3).

**Fig. 6.**
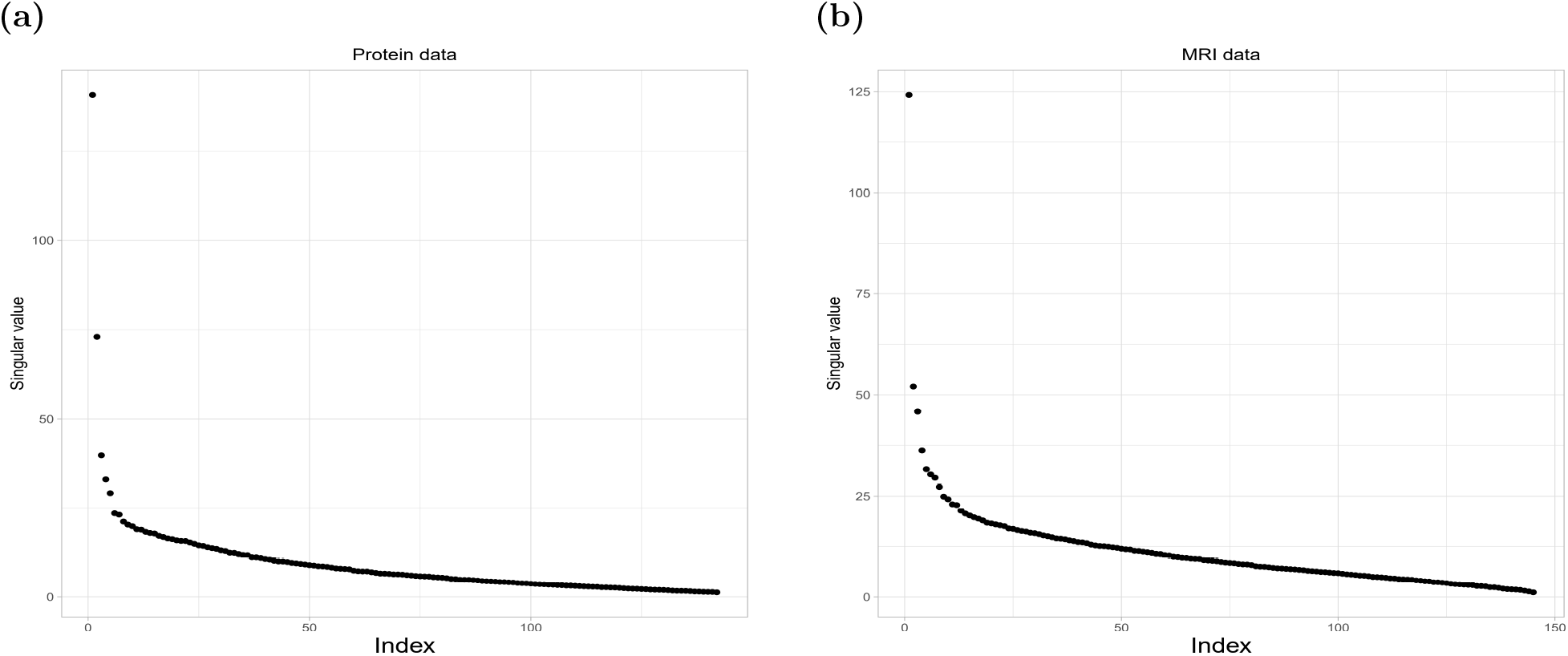
Singular values of standardized exposure and mediator data. (a) is corresponding to the protein data, while (b) is corresponding to the MRI data.

Following the preprocessing of data, we apply our method to the processed data. However, after implementing the BH procedure, no significant paths are obtained when controlling the FDR at a nominal level of 0.1. In order to obtain meaningful results, we relax the criterion and set the significance threshold for p-values to 0.1 without applying multiple correction. Consequently, we identify 63 significant causal paths, corresponding to 45 proteins and 7 brain regions. The estimated path effects, including the *θ*_*kl*_ and *β*_*l*_, are presented in the Supplementary Table S1. In Figure 7, we visualize the significant causal paths.

**Fig. 7.**
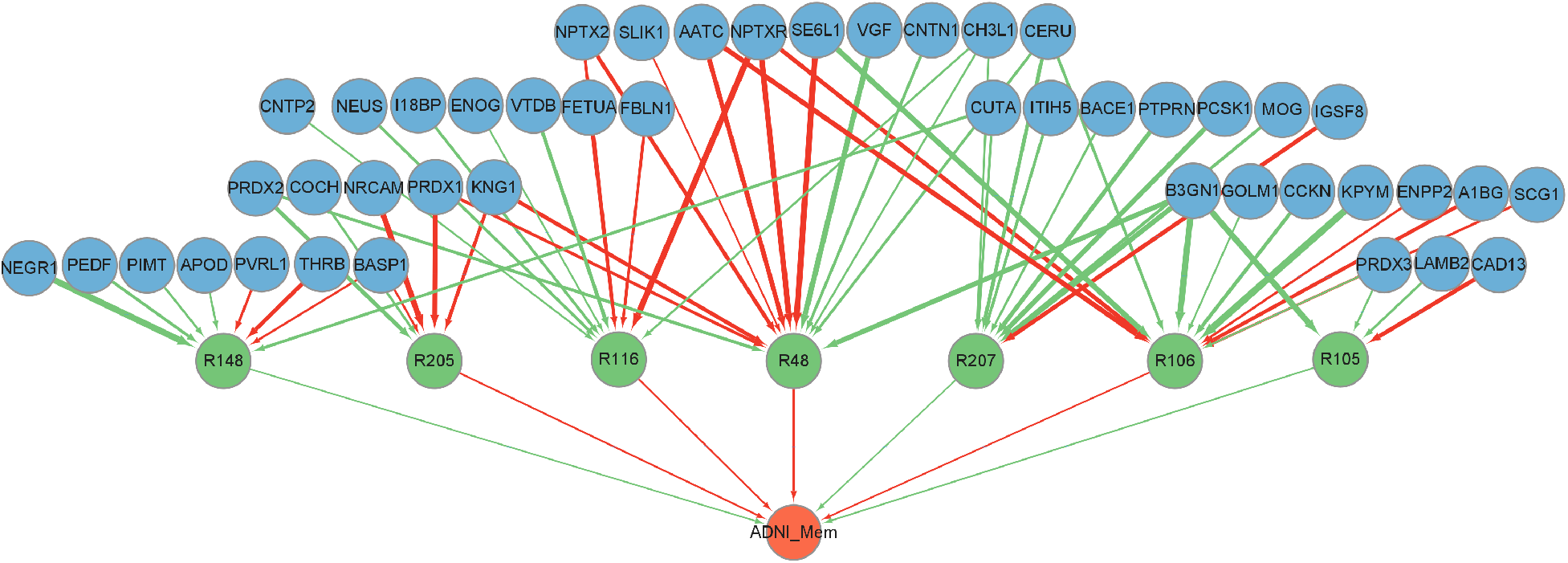
The estimated significant causal paths using proteomics-radiomics data. Blue nodes represent the proteins as exposures, green nodes represent the brain regions as mediators and the red node represents the memory score as the outcome. Red lines indicate positive estimates while green lines represent negative estimates. Line thickness corresponds to the effect size.

Our study has identified several brain regions associated with cognitive impairment and AD. Among them, R48 (left hippocampus) plays a crucial role in learning and memory, and is particularly vulnerable to early-stage damage in AD [Nadel and Hardt, 2011]. Importantly, hippocampal atrophy has been universally recognized and validated as the most reliable biomarker for AD [Schröder and Pantel, 2016]. Another crucial region in cognition is R106 (right angular gyrus), which is associated with language, spatial, and memory functions [Seghier, 2013, Humphreys et al., 2021]. The aging process leads to structural atrophy in the angular gyrus, which is linked to subjective and mild cognitive impairments, as well as dementia [Karas et al., 2008, Jockwitz et al., 2023]. Additionally, another significant region, R116 (right entorhinal area), often exhibits the earliest histological alterations in AD. Impaired neuronal activity in the area may cause memory impairments and spatial navigation deficits at the initial stage of AD [Igarashi, 2023]. Furthermore, R205 (left triangular part of the inferior frontal gyrus), R105 (left anterior orbital gyrus), R148 (right postcentral gyrus medial segment) and R207 (left transverse temporal gyrus) are also associated with AD and cognitive impairment. However, further investigation is necessary to comprehensively elucidate the roles of these regions in AD pathology and cognitive function.

Several proteins have been identified as potentially critical biomarkers for AD. NPTX2 and NPTXR are proteins that bind to glutamate receptors, contributing to synaptic plasticity. Reductions in NPTX2 have been linked to disruptions of the pyramidal neuron-PV interneuron circuit in an AD mouse model [Xiao et al., 2017]. PRDX1 and PRDX2 are peroxiredoxin proteins that provide protection against neuronal cell death and oxidative stress [Kim et al., 2001]. PRDX3 plays a crucial role as a mitochondrial antioxidant defense enzyme, and its overexpression provides protection against cognitive impairment while reducing the accumulation of A*β* in transgenic mice [Chen et al., 2012]. Furthermore, its overexpression reduces mitochondrial oxidative stress, attenuates memory impairment induced by hydrogen peroxide and improves cognitive ability in transgenic mice [Chen et al., 2014]. Moreover, recent research has revealed that PRDX3 plays important roles in neurite outgrowth and the development of AD [Xu et al., 2022a]. KNG1 is a protein involved in inflammatory responses, and leavage of KNG1 has been associated with the release of proinflammatory bradykininwhich may contribute to AD-associated inflammation [Markaki et al., 2020]. SE6L1 is a potential neuronal substrate of the AD protease BACE1, which is a major drug target in AD [Pigoni et al., 2016]. Aberrant function of SE6L1 may lead to movement disorders and neuropsychiatric diseases [Ong-Pålsson et al., 2022]. Overexpression of the neuropeptide precursor VGF has been found to partially rescue A*β* mediated memory impairment and neuropathology in a mouse model, indicating a protective function against the development and progression of AD [Beckmann et al., 2020]. CUTA is a protein that has been proposed to mediate acetylcholinesterase activity and copper homeostasis, which are important events in AD pathology. Overexpression of CUTA can reduce BACE1-mediated APP processing and A*β* generation, while RNA interference increases it [Zhao et al., 2012]. PEDF is a unique neurotrophic and neuroprotective protein whose expression decays with aging. Experiments in a senescence-accelerated mouse model show that PEDF negatively regulates A*β* and notably reduces cognitive impairment, suggesting that PEDF might play a crucial role in the development of AD [Huang et al., 2018]. Knock-down of PIMT and treatment with AdOX significantly increase A*β* secretion, which serves as a negative regulator of A*β* peptide formation and a potential protective factor in the pathogenesis of AD [Bae et al., 2011].

In summary, our study identified several critical brain regions, such as R48, R106 and R116, that are associated with learning, memory, and recognition. Moreover, we have identified several potential biomarkers for AD, such as NPTX2, NPTXR, PRDX1, PRDX2, PRDX3, KNG1, SE6L1, VGF, CUTA, PEDF, etc., most of which are not selected by the method ZY [Zhao et al., 2022]. Nonetheless, it is crucial to note that these findings are only suggestive and further experimental validation is warranted to fully understand their contributions to AD pathology and cognitive function.

## 5 Discussion

In this paper, we propose HILAMA, a new method for high-dimensional mediation analysis, an important statistical task in the analysis of multi-omics datasets increasingly available in biomedical sciences. HILAMA effectively unravels the causal pathway between high-dimensional exposures and a continuous outcome, in the presence of possibly latent/unmeasured confounders. We validate the practical performance of HILAMA through extensive simulations and by applying it to a real ADNI dataset, which allows for the identification of potential biomarkers for Alzheimer’s disease.

HILAMA features several key advantages over previous methods, designed towards better fitting into real-world multi-omics datasets. First, it is the first method to consider both high-dimensional exposures and high-dimensional mediators in the presence of latent confounders without transforming exposures/mediators into principal components, rendering the analysis results more interpretable. Second, it incorporates a novel Decorrelate & Debias method [Sun et al., 2022] to handle latent/unmeasured confounding and improve coefficient estimation, leading to better FDR control. Third, it employs a MinScreen screening procedure [Djordjilovi et al., 2019] to reduce the number of hypotheses being tested, thereby enhancing the statistical power of the tests. Finally, the method is computationally efficient and has implemented parallel computing techniques to handle the ever-increasing size and dimension of modern multi-omics datasets.

To conclude, we point out several venues for future research. First, HILAMA assumes linear models, which is standard practice in multi-omics studies. However, it will be interesting to generalize it to nonlinear/nonparametric models via nonlinear factor analysis [Amemiya and Yalcin, 2001, Feng, 2020], autoencoders [Yang et al., 2021], kernel methods [Singh et al., 2021] or deep neural networks [Xu et al., 2022b]. Second, HILAMA assumes that the effects of latent/unmeasured confounders on observables are dense. It may be possible to relax this assumption by extending the randomized data-augmentation scheme proposed in Zhang et al. [2022b] for total effect to the mediation analysis setting. Third, considering the correlation structure among mediators could be beneficial in the step of Mediator-Exposure deconfounded regression. Currently, this correlation structure is omitted due to challenges in controlling FDR in large-scale regression with multivariate response. However, incorporating this information might be useful for removing the confounding effect and improving the power of the test [Zou et al., 2020, Kotekal and Gao, 2021]. Finally, other methods of dealing with latent confounding can also be incorporated into HILAMA in its future version, such as the approaches [Miao et al., 2022, Tang et al., 2023] that directly leverage the majority rule [Kang et al., 2016] or the plurality rule [Guo et al., 2018]. Overall, these future research directions have the potential to expand the capabilities of HILAMA, allowing for more accurate and robust causal inference in multi-omics studies.

## Acknowledgments

The authors would like to thanks professor Yi Zhao for her valuable suggestions on accessing the ADNI data.

Data collection and sharing for this project was funded by the Alzheimer’s Disease Neuroimaging Initiative (ADNI) (National Institutes of Health Grant U01 AG024904) and DOD ADNI (Department of Defense award number W81XWH-12-2-0012). ADNI is funded by the National Institute on Aging, the National Institute of Biomedical Imaging and Bioengineering, and through generous contributions from the following: AbbVie, Alzheimer’s Association; Alzheimer’s Drug Discovery Foundation; Araclon Biotech; BioClinica, Inc.; Biogen; Bristol-Myers Squibb Company; CereSpir, Inc.; Cogstate; Eisai Inc.; Elan Pharmaceuticals, Inc.; Eli Lilly and Company; EuroImmun; F. Hoffmann-La Roche Ltd and its affiliated company Genentech, Inc.; Fujirebio; GE Healthcare; IXICO Ltd.; Janssen Alzheimer Immunotherapy Research & Development, LLC.; Johnson & Johnson Pharmaceutical Research & Development LLC.; Lumosity; Lundbeck; Merck & Co., Inc.; Meso Scale Diagnostics, LLC.; NeuroRx Research; Neurotrack Technologies; Novartis Pharmaceuticals Corporation; Pfizer Inc.; Piramal Imaging; Servier; Takeda Pharmaceutical Company; and Transition Therapeutics. The Canadian Institutes of Health Research is providing funds to support ADNI clinical sites in Canada. Private sector contributions are facilitated by the Foundation for the National Institutes of Health (www.fnih.org). The grantee organization is the Northern California Institute for Research and Education, and the study is coordinated by the Alzheimer’s Therapeutic Research Institute at the University of Southern California. ADNI data are disseminated by the Laboratory for Neuro Imaging at the University of Southern California.

## Supplementary Materials

### Derivations of direct and indirect effects under LSEM (1) and (2)

We follow the potential outcomes framework [Rubin, 1972, Splawa-Neyman et al., 1990] to decompose the effect of exposures on the outcome. Specifically, we denote *Y*(***x, m***) as the potential outcome when the exposure ***X*** is set to ***x*** = (*x*_1_, ⋯, *x*_*p*_)′, the mediator ***M*** is set to ***m*** = (*m*_1_, ⋯, *m*_*q*_)′. Formally, we define ***x***_−*k*_ = (*x*_1_, ⋯, *x*_*k*−1_, *x*_*k*+1_, ⋯, *x*_*p*_)′ and use 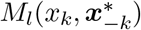 as the potential value of the *l*-th mediator where the *k*-th exposure is set to *x*_*k*_ and the ot her exposures are set to 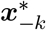. We denote the baseline adjusted covariates as ***Z*** = (***Z***_1_, ***H***), where ***Z***_1_ represents the observ ed confounders, ***H*** represents the potential latent confounders, and as the reference level of exposure. Then, the average total effect (TE) of the *k*-th exposure on the outcome is defined as 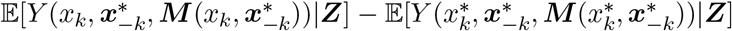 the average natural direct effect (NDE) of*X*_*k*_ on *Y* is 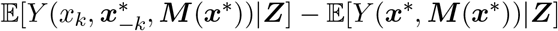 the average natural indirect effect (NIE) of *X*_*k*_ on *Y* is 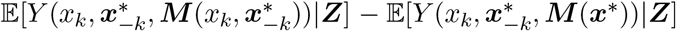 Among the three effects, we have the following relationships:

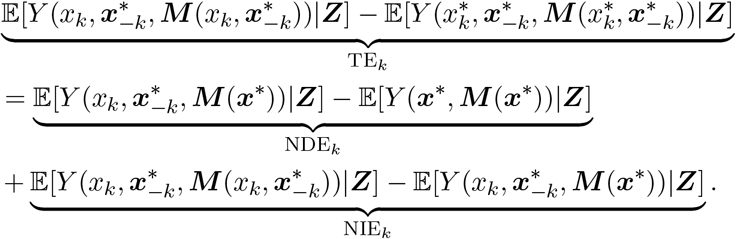

To identify the above NDE and NIE, we need the following standard ignorability assumption [VanderWeele and Vansteelandt, 2014]:

(C1) *Y*(***x, m***) ⊥ *X*_*k*_| ***Z*** for ∀***x, m***, *k* ∈ [*p*], i.e. no unmeasured confounding between the exposures and the outcome;

(C2) *Y*(***x, m***) ⊥ *M*_*l*_ | ***Z*** for ∀***x, m***, *l* ∈ [*q*], i.e. no unmeasured confounding between the mediators and the outcome;

(C3) *M*_*l*_(***x***) ⊥ *X*_*k*_ | ***Z*** for ∀ ***x***, ∈ *k* [*p*], *l* ∈ [*q*], i.e. no unmeasured confounding between the exposures and the outcome;

(C4) *Y*(***x, m***) ⊥ *M*_*l*_(***x***\*) |***Z*** for ∀***x, x***\*, ***m***, *l* ∈ [*q*], i.e. no unmeasured confounding between the mediators and the outcome that is itself affected by the exposures.

To simplify the representation, we replace ***H*** in LSEM (1) and (2) with ***Z*** to represent the above baseline covariates ***Z*** = (***Z***_1_, ***H***). From LSEM (2), we can express the potential mediator 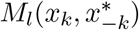 and potential outcome 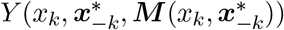 as follows:

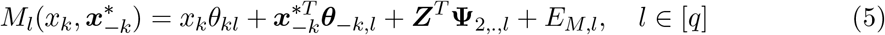

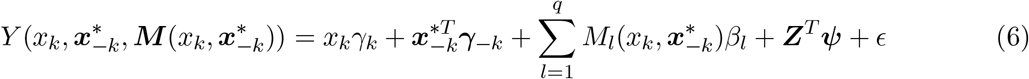

Then, the (average) natural direct effect of exposure *X*_*k*_ on outcome when the value of that exposure is manipulated from *x*_*k*_ to 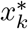 denoted by 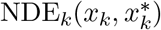 can be derived directly based on equation (6):

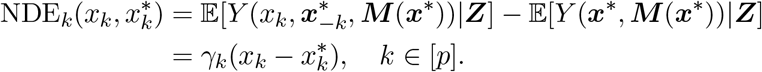

Similarly, the natural indirect effect of exposure *X*_*k*_ on outcome, denoted by 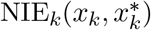 can be derived as follows:

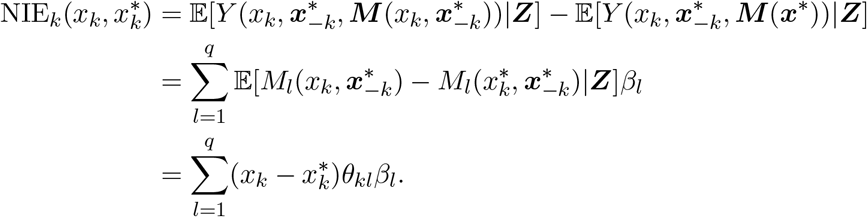

## Supplementary tables and figures

**Table S1.**
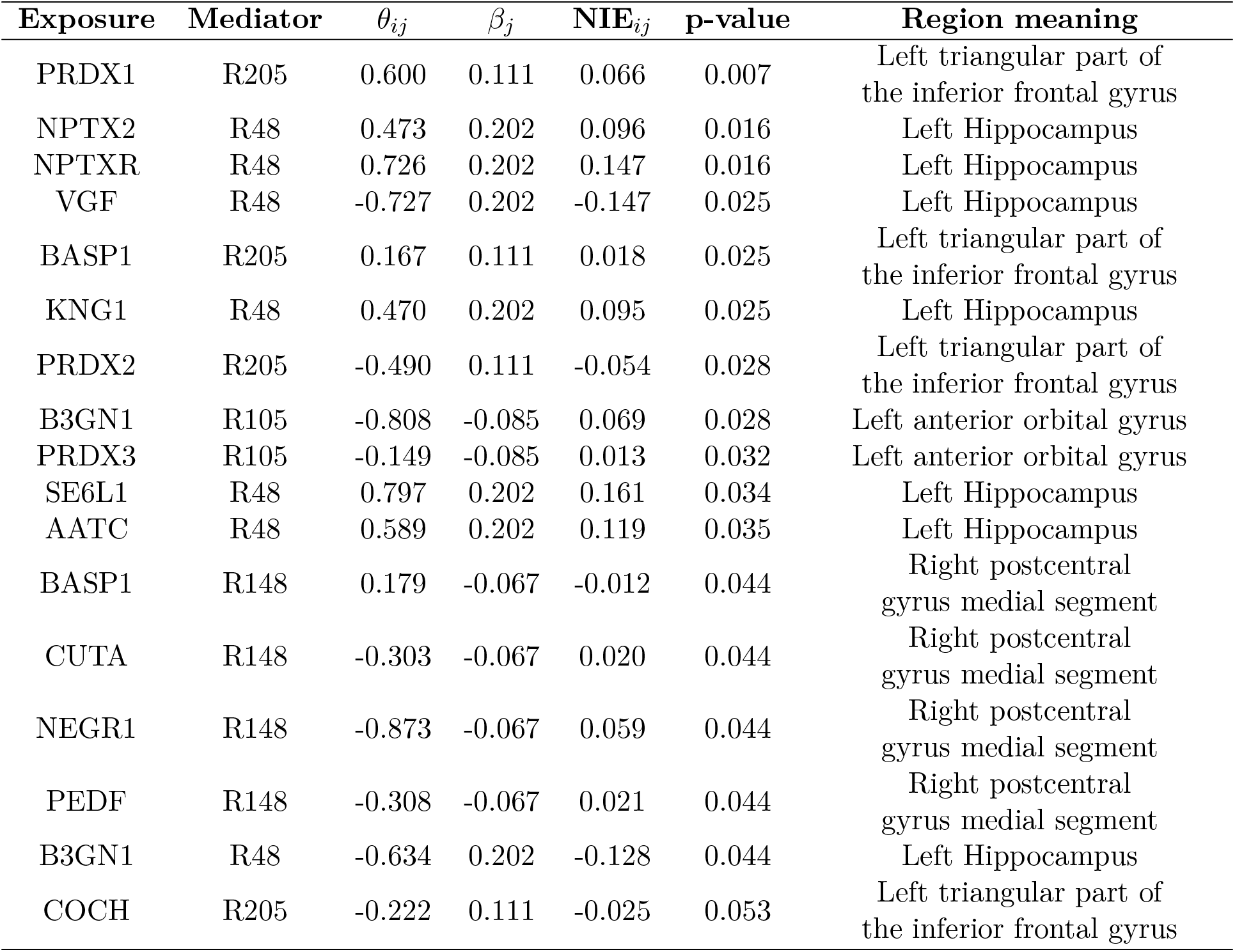
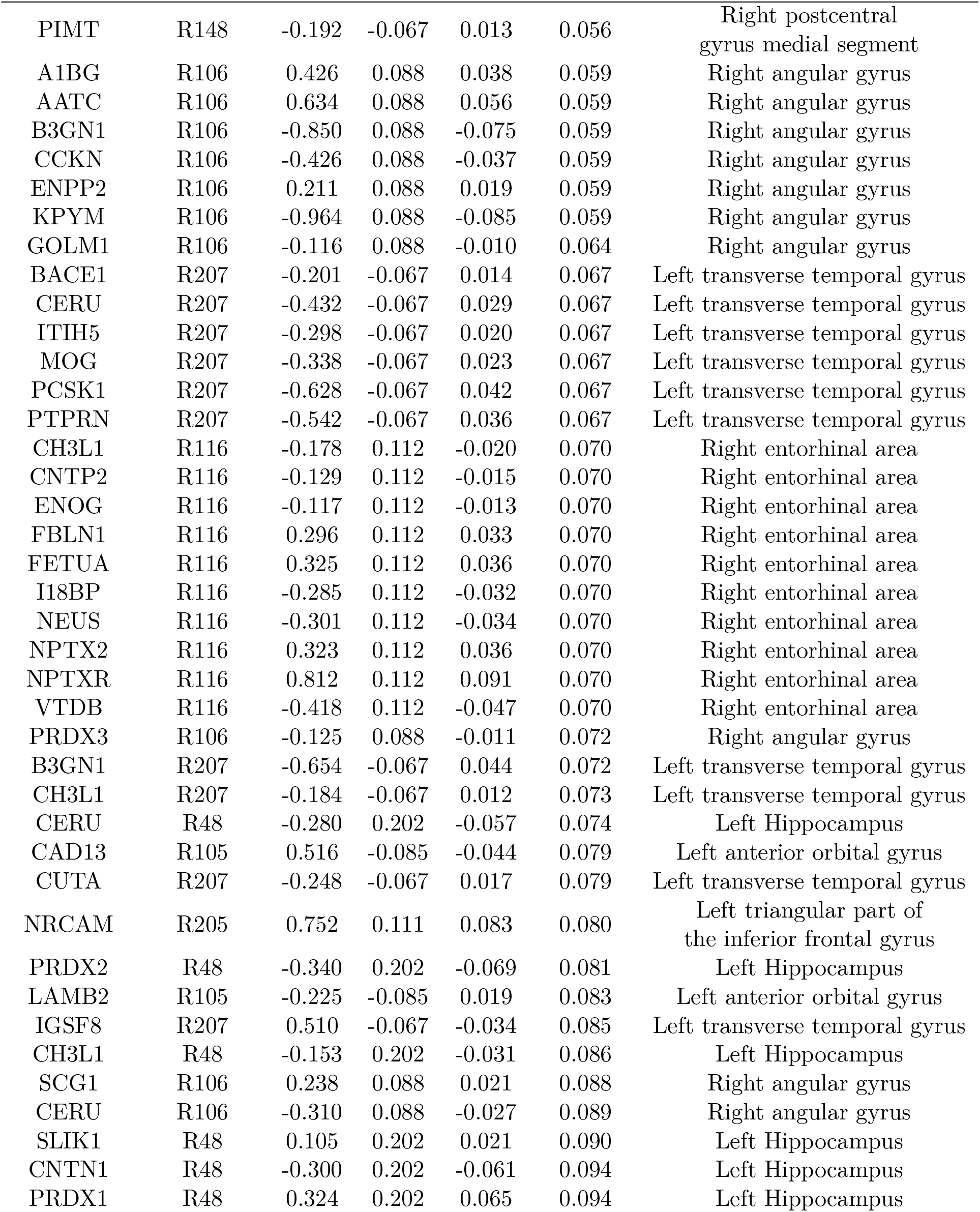
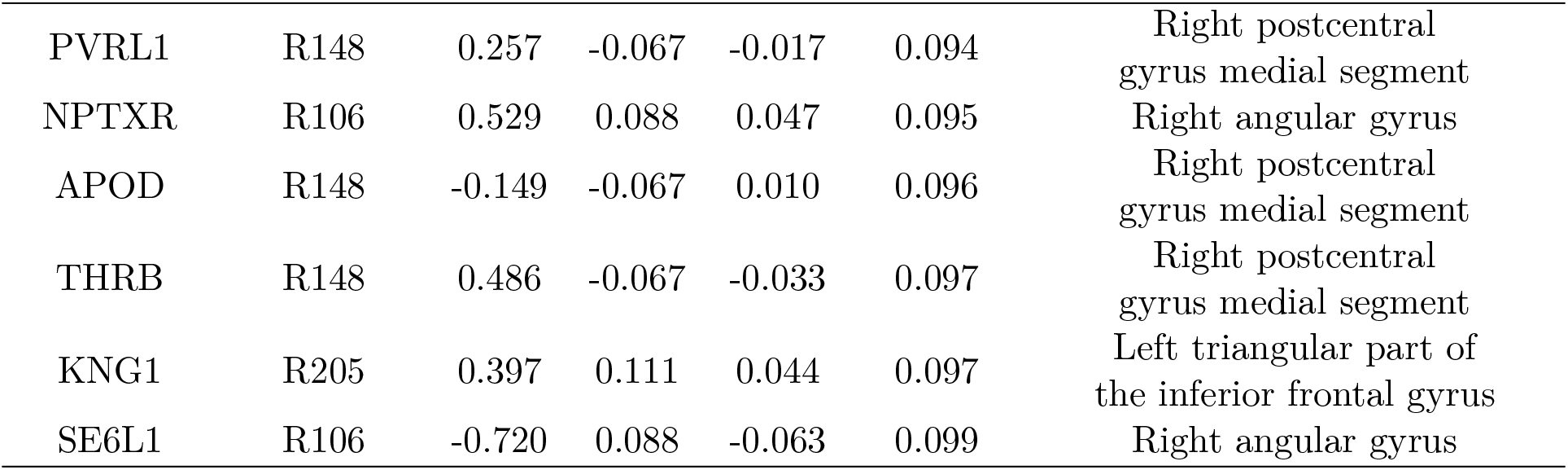
The summary statistics of the estimated significant paths.

| Exposure | Mediator | $\theta_{ij}$ | $\beta_j$ | $\text{NIE}_{ij}$ | p-value | Region meaning |
| --- | --- | --- | --- | --- | --- | --- |
| PRDX1 | R205 | 0.600 | 0.111 | 0.066 | 0.007 | Left triangular part of the inferior frontal gyrus |
| NPTX2 | R48 | 0.473 | 0.202 | 0.096 | 0.016 | Left Hippocampus |
| NPTXR | R48 | 0.726 | 0.202 | 0.147 | 0.016 | Left Hippocampus |
| VGF | R48 | -0.727 | 0.202 | -0.147 | 0.025 | Left Hippocampus |
| BASP1 | R205 | 0.167 | 0.111 | 0.018 | 0.025 | Left triangular part of the inferior frontal gyrus |
| KNG1 | R48 | 0.470 | 0.202 | 0.095 | 0.025 | Left Hippocampus |
| PRDX2 | R205 | -0.490 | 0.111 | -0.054 | 0.028 | Left triangular part of the inferior frontal gyrus |
| B3GN1 | R105 | -0.808 | -0.085 | 0.069 | 0.028 | Left anterior orbital gyrus |
| PRDX3 | R105 | -0.149 | -0.085 | 0.013 | 0.032 | Left anterior orbital gyrus |
| SE6L1 | R48 | 0.797 | 0.202 | 0.161 | 0.034 | Left Hippocampus |
| AATC | R48 | 0.589 | 0.202 | 0.119 | 0.035 | Left Hippocampus |
| BASP1 | R148 | 0.179 | -0.067 | -0.012 | 0.044 | Right postcentral gyrus medial segment |
| CUTA | R148 | -0.303 | -0.067 | 0.020 | 0.044 | Right postcentral gyrus medial segment |
| NEGR1 | R148 | -0.873 | -0.067 | 0.059 | 0.044 | Right postcentral gyrus medial segment |
| PEDF | R148 | -0.308 | -0.067 | 0.021 | 0.044 | Right postcentral gyrus medial segment |
| B3GN1 | R48 | -0.634 | 0.202 | -0.128 | 0.044 | Left Hippocampus |
| COCH | R205 | -0.222 | 0.111 | -0.025 | 0.053 | Left triangular part of the inferior frontal gyrus |

| Exposure | Mediator | $\theta_{ij}$ | $\beta_j$ | NIE$_{ij}$ | p-value | Region meaning |
| --- | --- | --- | --- | --- | --- | --- |
| PIMT | R148 | -0.192 | -0.067 | 0.013 | 0.056 | Right postcentral gyrus medial segment |
| A1BG | R106 | 0.426 | 0.088 | 0.038 | 0.059 | Right angular gyrus |
| AATC | R106 | 0.634 | 0.088 | 0.056 | 0.059 | Right angular gyrus |
| B3GN1 | R106 | -0.850 | 0.088 | -0.075 | 0.059 | Right angular gyrus |
| CCKN | R106 | -0.426 | 0.088 | -0.037 | 0.059 | Right angular gyrus |
| ENPP2 | R106 | 0.211 | 0.088 | 0.019 | 0.059 | Right angular gyrus |
| KPYM | R106 | -0.964 | 0.088 | -0.085 | 0.059 | Right angular gyrus |
| GOLM1 | R106 | -0.116 | 0.088 | -0.010 | 0.064 | Right angular gyrus |
| BACE1 | R207 | -0.201 | -0.067 | 0.014 | 0.067 | Left transverse temporal gyrus |
| CERU | R207 | -0.432 | -0.067 | 0.029 | 0.067 | Left transverse temporal gyrus |
| ITIH5 | R207 | -0.298 | -0.067 | 0.020 | 0.067 | Left transverse temporal gyrus |
| MOG | R207 | -0.338 | -0.067 | 0.023 | 0.067 | Left transverse temporal gyrus |
| PCSK1 | R207 | -0.628 | -0.067 | 0.042 | 0.067 | Left transverse temporal gyrus |
| PTPRN | R207 | -0.542 | -0.067 | 0.036 | 0.067 | Left transverse temporal gyrus |
| CH3L1 | R116 | -0.178 | 0.112 | -0.020 | 0.070 | Right entorhinal area |
| CNTP2 | R116 | -0.129 | 0.112 | -0.015 | 0.070 | Right entorhinal area |
| ENOG | R116 | -0.117 | 0.112 | -0.013 | 0.070 | Right entorhinal area |
| FBLN1 | R116 | 0.296 | 0.112 | 0.033 | 0.070 | Right entorhinal area |
| FETUA | R116 | 0.325 | 0.112 | 0.036 | 0.070 | Right entorhinal area |
| I18BP | R116 | -0.285 | 0.112 | -0.032 | 0.070 | Right entorhinal area |
| NEUS | R116 | -0.301 | 0.112 | -0.034 | 0.070 | Right entorhinal area |
| NPTX2 | R116 | 0.323 | 0.112 | 0.036 | 0.070 | Right entorhinal area |
| NPTXR | R116 | 0.812 | 0.112 | 0.091 | 0.070 | Right entorhinal area |
| VTDB | R116 | -0.418 | 0.112 | -0.047 | 0.070 | Right entorhinal area |
| PRDX3 | R106 | -0.125 | 0.088 | -0.011 | 0.072 | Right angular gyrus |
| B3GN1 | R207 | -0.654 | -0.067 | 0.044 | 0.072 | Left transverse temporal gyrus |
| CH3L1 | R207 | -0.184 | -0.067 | 0.012 | 0.073 | Left transverse temporal gyrus |
| CERU | R48 | -0.280 | 0.202 | -0.057 | 0.074 | Left Hippocampus |
| CAD13 | R105 | 0.516 | -0.085 | -0.044 | 0.079 | Left anterior orbital gyrus |
| CUTA | R207 | -0.248 | -0.067 | 0.017 | 0.079 | Left transverse temporal gyrus |
| NRCAM | R205 | 0.752 | 0.111 | 0.083 | 0.080 | Left triangular part of the inferior frontal gyrus |
| PRDX2 | R48 | -0.340 | 0.202 | -0.069 | 0.081 | Left Hippocampus |
| LAMB2 | R105 | -0.225 | -0.085 | 0.019 | 0.083 | Left anterior orbital gyrus |
| IGSF8 | R207 | 0.510 | -0.067 | -0.034 | 0.085 | Left transverse temporal gyrus |
| CH3L1 | R48 | -0.153 | 0.202 | -0.031 | 0.086 | Left Hippocampus |
| SCG1 | R106 | 0.238 | 0.088 | 0.021 | 0.088 | Right angular gyrus |
| CERU | R106 | -0.310 | 0.088 | -0.027 | 0.089 | Right angular gyrus |
| SLIK1 | R48 | 0.105 | 0.202 | 0.021 | 0.090 | Left Hippocampus |
| CNTN1 | R48 | -0.300 | 0.202 | -0.061 | 0.094 | Left Hippocampus |
| PRDX1 | R48 | 0.324 | 0.202 | 0.065 | 0.094 | Left Hippocampus |
| PVRL1 | R148 | 0.257 | -0.067 | -0.017 | 0.094 | Right postcentral gyrus medial segment |
| NPTXR | R106 | 0.529 | 0.088 | 0.047 | 0.095 | Right angular gyrus |
| APOD | R148 | -0.149 | -0.067 | 0.010 | 0.096 | Right postcentral gyrus medial segment |
| THRB | R148 | 0.486 | -0.067 | -0.033 | 0.097 | Right postcentral gyrus medial segment |
| KNG1 | R205 | 0.397 | 0.111 | 0.044 | 0.097 | Left triangular part of the inferior frontal gyrus |
| SE6L1 | R106 | -0.720 | 0.088 | -0.063 | 0.099 | Right angular gyrus |

**Fig. S1.**
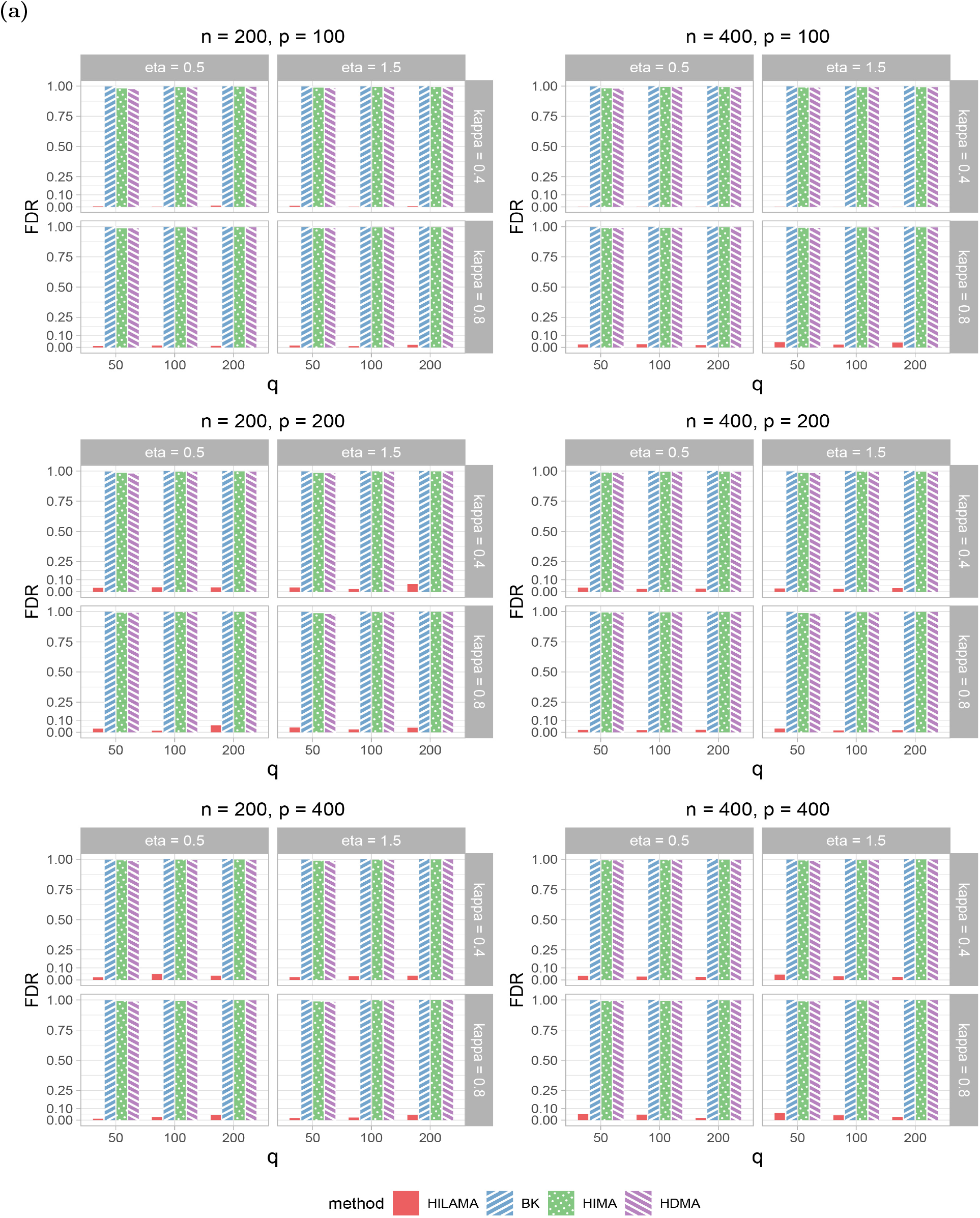
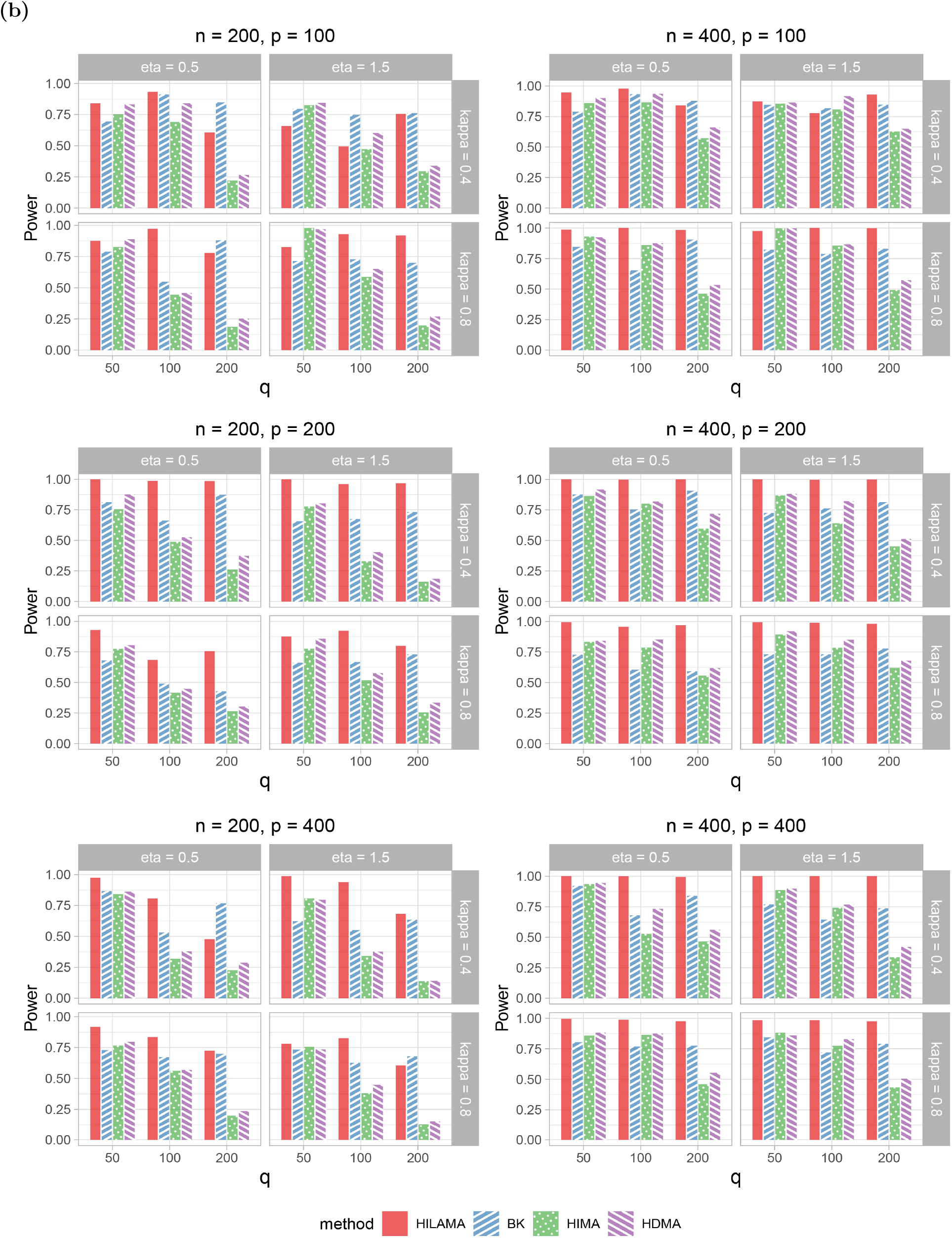
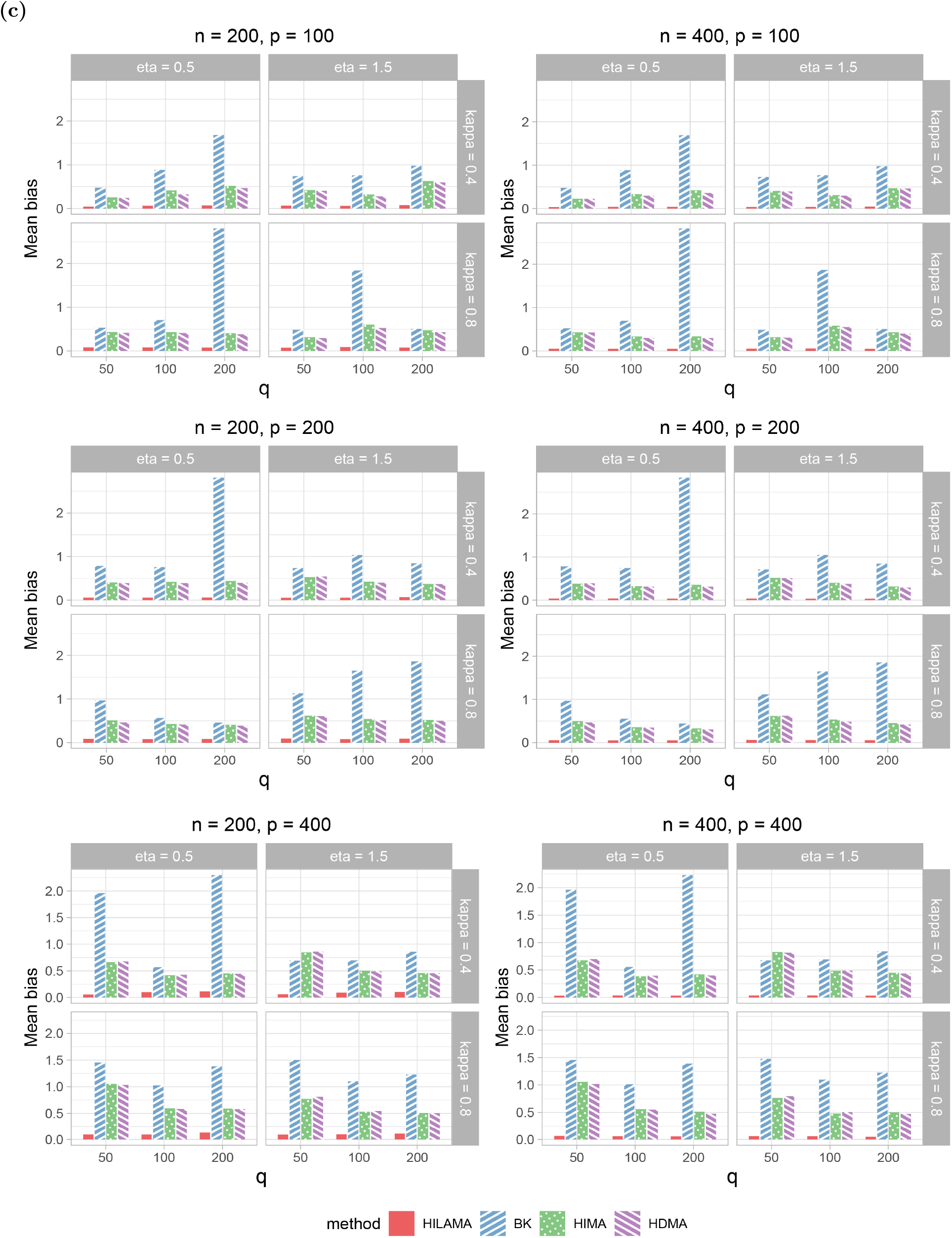
Comparison results of (a) Empirical False Discovery Rate (FDR), (b) Empirical Power and (c) Empirical Mean bias for different methods in **simulation 1** across 72 scenarios. *eta* represents latent effect and *kappa* represents the correlation size among exposure. All the results are averaged over 50 replications under the nominal FDR level of 0.1.

**Fig. S2.**
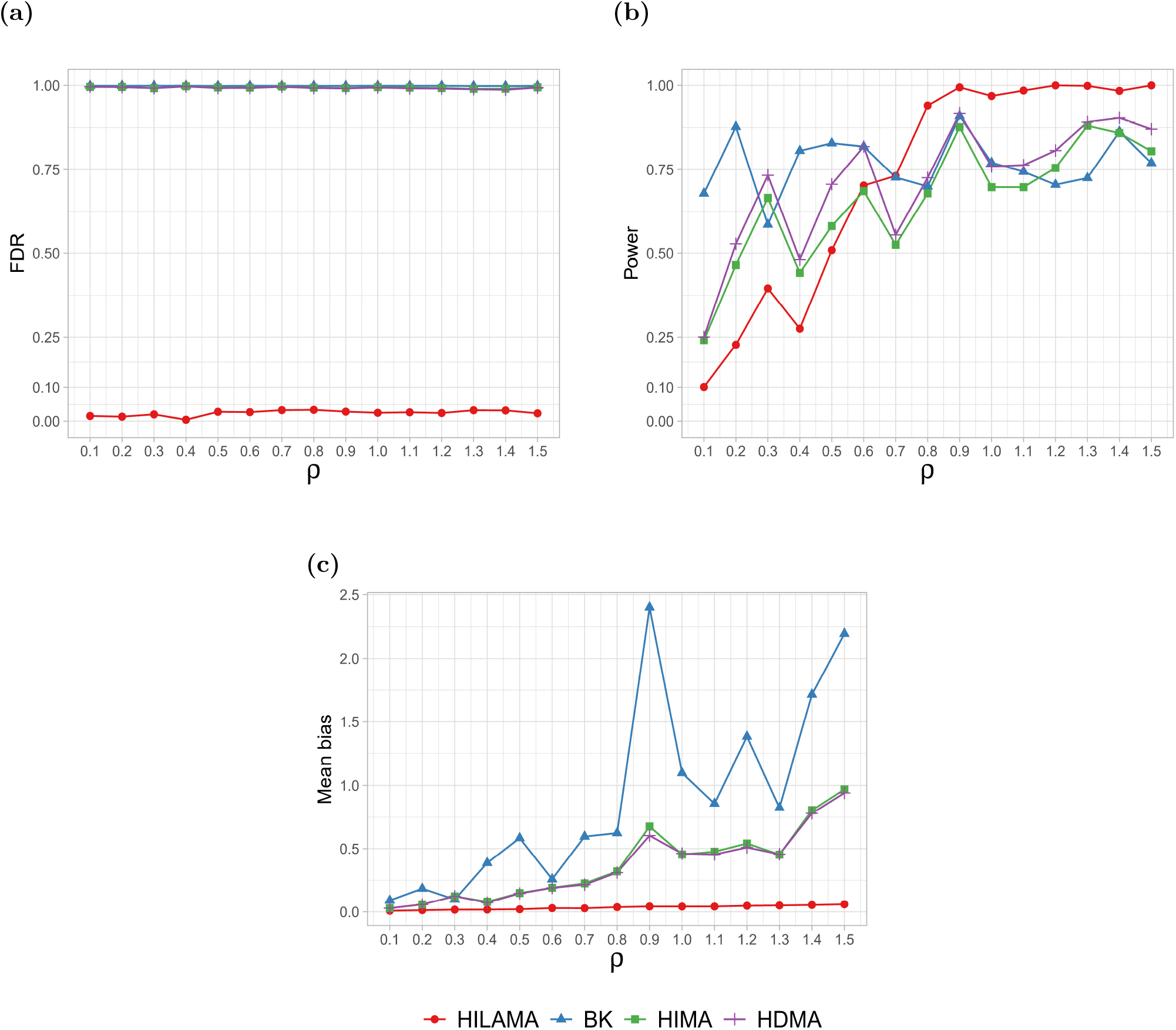
Comparison results of (a) Empirical FDR, (b) Empirical Power and (c) Mean bias for different methods in **simulation 3** with varied signal strength *ρ* while fixing *n* = 400, *p* = 300, *q* = 100, *κ* = 0.6, *η* = 1, *r*_*p*_ = *r*_*pq*_ = 0.1, *s* = 3. All results are averaged over 50 replications under the nominal FDR level of 0.1.

